# CCIDeconv: Hierarchical model for deconvolution of subcellular cell-cell interactions in single-cell data

**DOI:** 10.64898/2026.03.26.714643

**Authors:** Rojashree Jayakumar, Pratibha Panwar, Jean Yee Hwa Yang, Shila Ghazanfar

## Abstract

Cell-cell interaction (CCI) underlies several fundamental biological processes, including development, homeostasis and disease progression. Subcellular spatial transcriptomics (sST) provides an opportunity to examine whether CCI-associated signals show compartment-specific patterns within cells. Assessing CCI at subcellular level can help us gain insights into the distinct pathway activation and signalling patterns. We developed a novel approach that deconvolutes CCI into subcellular CCI (sCCI) information from non-spatial single-cell transcriptomics (scRNA-seq) based CCI using a modified CellChat-derived communication score. By estimating communication scores separately for cytoplasmic and nuclear compartments, we identified compartment-associated sCCI. We then deconvolved whole-cell communication scores into subcellular compartments using a hierarchical classification and regression framework, which we call CCIDeconv. To ensure biological fidelity, we integrated protein localisation data from the Human Protein Atlas in our deconvolution model. Across nine publicly available human sST datasets, leave-one-dataset-out validation achieved a median composite metric of 0.75, with mean *R*^2^ values of 0.87 and 0.80 for cytoplasmic- and nuclear-associated scores, respectively. Performance without spatial features approached that of spatial models as the number of training datasets increased, supporting application to non-spatial scRNA-seq data. This highlighted the potential for prediction of sCCI from scRNA-seq, given a suffciently large number of training datasets. Overall, our method can attribute whole-cell CCI to its subcellular compartments, allowing researchers to dissect sCCI patterns and gain insights into the underlying biology of healthy and disease tissues.

## Introduction

Cell-cell interaction (CCI) is important in understanding biological systems and occurs through intracellular and intercellular mechanisms, involving signal generation, transmission, and subsequent signal transduction (Su et al., 2024). These events are mediated by ligands on the signaling cell and receptors on the target cell. Ligand and receptors (LR) can range from extracellular proteins to secreted molecules such as ions, cytokines and hormones. Therefore, cells communicate with each other using a diverse range of molecules and signal various downstream pathways that result in altered gene expression (Armingol et al., 2020). Advances in single-cell transcriptomics have enabled systematic investigation of CCI through computational inference.

With a wealth of high-throughput single-cell transcriptomics data, there have been tremendous efforts in the inference of CCI at the individual cell and cell type levels. Given a database of LR pairs and complexes, e.g. OmniPath (Türei et al., 2016), CellPhoneDB (Troulé et al., 2025), and CellChatDB (Jin et al., 2021), and a single-cell dataset, methods such as CellChat, NicheNet (Browaeys et al., 2020), and CellPhoneDB identify CCI that occur in the given tissue. Methods such as MISTy (Tanevski et al., 2022) and NicheCompass (Birk et al., 2025) have been developed to utilise the additional spatial coordinates of cells to perform inference of CCI. Similarly, methods originally developed for single-cell dissociated data have undergone additional development to account for spatial coordinates, including CellChat (Jin et al., 2025) and CellPhoneDB (v5). Some methods such as DeepLinc (Li and Yang, 2022) identify de novo CCI without the need for a curated database. CellChat calculates the communication score as a function of the expression and assesses significance by permutation testing, while other methods such as NicheNet use network based approaches to calculate the scores.

Despite these advances, existing methods infer communication at the cellular level and assume that ligand and receptor expression is homogeneous within individual cells. This assumption can be revisited with the emergence of subcellular spatial transcriptomics (sST) technologies, which provide quantification of gene expression at intracellular resolution and typically rely on microscopy imaging. sST may also be sequencing-based that offer subcellular resolution through DNA nanoballs (Stereo-seq (Zhao et al., 2025)) or extremely high resolution chips (10X Genomics Visium HD (Nagendran et al., 2023)). Current CCI methods leverage spatial relationships between cells but do not exploit the additional subcellular information available within cells. As a result, potentially important compartment-specific signalling patterns are diluted or entirely masked in whole-cell averages.

Subcellular CCI (sCCI) can determine where CCI is initiated, and which signaling pathways are activated, influence disease mechanisms, or guide therapeutic targeting. For example, GPCRs can initiate signalling in both the plasma membrane and intracellular organelles, producing distinct downstream effects and cellular functions (Bock et al., 2025). Similarly, engaging serotonin 2A receptors inside neurons promotes neuronal growth, whereas activation of the same receptor at the cell surface does not (Vargas et al., 2023). Extracellular vesicle-mediated delivery of ligands provides further evidence that CCI occur within intracellular compartments (Lorico et al., 2025). Some interactions occur on the surface; for example, microglial activation is based on LR recognition on the cell surface in Alzheimer’s disease (Chen et al., 2006) while others such as specific tumor-stromal signaling cascades occur inside the cytoplasm (Gardiner and Cukierman, 2022). Together, these findings suggest that the subcellular location of ligand-receptor activity contains biologically meaningful information that is not captured by current cell-level CCI frameworks.

Here, we present CCIDeconv, a novel framework that uses sST data to deconvolve whole-cell LR communication scores into nuclear- and cytoplasmic-associated components. By integrating protein localisation annotations from the Human Protein Atlas (HPA) and CellChatDB as an additional source of biological evidence, CCIDeconv provides hypotheses about the likely subcellular context of inferred communication signals.

## Materials and methods

### Calculation of CellChat based communication score

We use LR pairs from the CellChatDB.human database to calculate the communication scores. We modified the CellChat formula by setting the agonist and antagonist expression components to a value of 1. This was necessary because the Xenium Prime 5K gene panel did not cover the agonist and antagonist genes defined in CellChatDB. Furthermore, we observed a high correlation between the communication scores calculated with and without these components (Supplementary Fig. 1).

**Figure 1.**
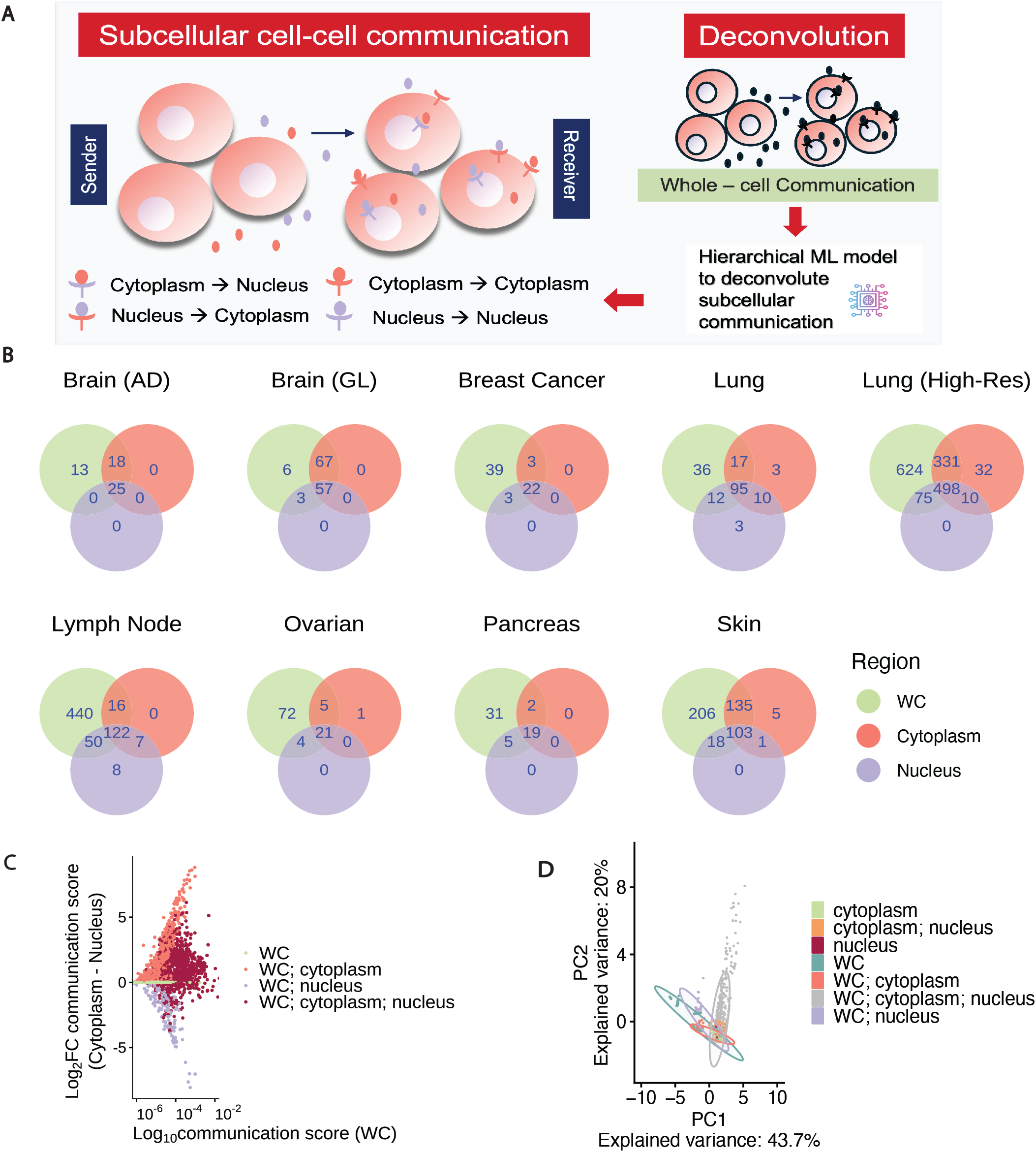
Differences in subcellular communication across nine different datasets. A. Overview of subcellular and whole-cell communication, and quantification of communication scores across the subcellular compartments. B. Significant interactions detected using CellChat are different between the cytoplasm and nucleus across the nine datasets. C. *Log*_2_ Fold Change differences in the LR communication score detected between the whole-cell and its subcellular compartments.D. PCA of CellChat-derived communication score components. WC - Whole-c ell

#### Communication score with spatial information

We calculate the whole-cell communication score as:

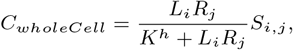

where *L*_*i*_ is the expression level of a ligand in cell type *i, R*_*j*_ is the expression level of a receptor in cell type *j*, and *S*_*i,j*_ is the spatial distance between cell types *i* and *j*. Also, *K* is the dissociation coeffcient and *h* is the Hill’s constant (Gesztelyi et al., 2012). Like in CellChat, *R*_*j*_ is weighted by expression of co-stimulatory (*RA*) and co-inhibitory receptors (*RI*) in cell type *j* and is calculated as 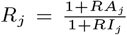 . We calculate *S*_*i,j*_ using the `computeRegionDistance()` function from CellChat. This function measures the average Euclidean distance between the nearest neighbours in celltype *i* and celltype *j*. We set *K* = 0.5 and *h* = 1 for all our scores as used by CellChat. Additionally, we define subcellular-specific CCI scores for the cytoplasm-cytoplasm and nucleus-nucleus communication as:

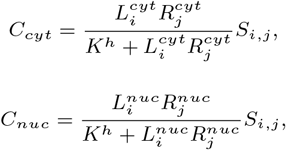

where 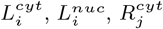 and 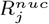 is the ligand and receptor expression level in the corresponding subcellular compartments.

#### Communication score without spatial information

For deconvolution of single-cell data, we set the spatial distance *S*_*i,j*_ to 1 when calculating the communication score, to effectively suppress the spatial component. It is calculated as

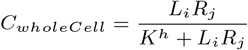

### Overview of CCIDeconv

#### CCIDeconv architecture

CCIDeconv uses a two-stage hierarchical framework. A classifier first determines whether a whole-cell CCI event contains suffcient compartment-specific signal for deconvolution. Two regression models then estimate its nuclear- and cytoplasmic-associated communication scores. We use the classifier to categorise each CCI event (ligand, receptor, sender cell types, and receiver cell types) detected in the whole-cell into a terminal class ***o*** or a separable class ***x*** . Class ***o*** represents CCI events that have low expression signal in subcellular compartments and cannot be deconvoluted. Class ***x*** represents CCI events whose communication scores can be deconvoluted. Finally, we use the two regression models on the CCI events in class ***x*** to obtain two communication scores, attributing to the nucleus and cytoplasm communication. We implemented two procedures using our CCIDeconv architecture:

1. Spatial procedure (SP): here we include *S*_*i,j*_ in the communication score and use it as one of the input features mentioned below in Feature scaling and encoding section.
2. Single-cell procedure (ScP): in this procedure, we do not include *S*_*i,j*_ in the communication score or as a feature.

Overall, CCIDeonv uses whole-cell communication score for each CCI event, alongside a suite of additional features (detailed in “Feature scaling and encoding section”), as the primary input. The model deconvolutes the whole-cell communication score into two distinct outputs for each CCI event, i.e., predicted communication scores for the cytoplasm and the nucleus for each specific CCI event.

#### Feature scaling and encoding

CCIDeconv input features are communication scores in the whole-cell, Hill function of the LR expression, receiver and sender cell types, ligand and receptor HGNC symbols as well as subcellular location and molecular information of ligands from HPA (The Human Protein Atlas; Uhlén et al., 2015) and CellChatDB. We assign a categorical feature to each ligand and receptor, representing its validated location from HPA and CellChatDB as priors (e.g., nucleoplasm, plasma membrane, or cytoplasm), to guide the deconvolution process. We scale the continuous features to zero mean and unit variance using `StandardScaler()` function from the scikit-learn library (Pedregosa et al., 2011) in Python. We apply target encoding to categorical features (Pargent et al., 2022). We also transform the location features into a binary indicator matrix using the `MultiLabelBinarizer()` function from scikit-learn library. The encoding of location features is based on the model selection procedure which we detail in “Model selection from different baselearner configurations and encoding location features section”. We perform scaling on each dataset and encoding separately for the training, test, and validation datasets to prevent data leakage.

#### Model specification

Within CCIDeconv, we use a voting classifier that combines Random Forest and XGBoost for classification, and two separate XGBoost regressors for subcellular score prediction. We selected these models based on evaluation of the candidate approaches detailed in “Model selection from different baselearner configurations and encoding location features” section. We train the classifier using a Logarithmic Loss function (binary cross-entropy), which optimises the probability of correctly assigning CCI events to class ***o*** or ***x*** . For the regression models, we train using a squared error loss function (`objective: reg:squarederror`), which minimizes the mean squared error between the predicted and ground-truth cytoplasmic and nuclear communication scores.

#### Hyperparameter tuning

For all models, we optimised hyperparameters using a Bayesian optimisation method (Garnett, 2023) implemented *via* the BayesianOptimization package in Python. The parameters used for tuning and the search space are detailed in the Supplementary methods 1.1.

### Preprocessing datasets

We obtained nine publicly-available sST datasets from the 10X Xenium platform (Supplementary methods 1.2.1.) and imported them into R using the `readXenium()` function from the MoleculeExperiment package (Peters Couto et al., 2023). We computed SpatialExperiment objects for nucleus and cytoplasm by aggregating the data using `countMolecules()`, with `boundariesAssay =` “*cell*” to obtain whole-cell level counts and `boundariesAssay =` “*nucleus*” to obtain nuclear counts. We calculated the cytoplasmic counts as the difference between the counts in the whole-cell and nucleus. Thus, we obtained three SpatialExperiment objects: whole-cell, cytoplasm, and nucleus for each dataset. Furthermore, we used `addPerCellQCMetrics()` function from the scuttle package (McCarthy et al., 2016) to compute quality control metrics and removed cells with total counts ≤ 20 and expressed genes ≤ 10. We excluded the negative-control probes prior to downstream analysis. We also normalised the cells using `logNormCounts()` function from the scuttle package. We used corresponding reference datasets for cell type annotation (detailed in Supplementary methods 1.2.2.) using the `singleR()` function from singleR package (Aran et al., 2019) in R. We then followed the standard spatial CellChat workflow and calculated the communication score as described above in calculation of CellChat based communication score. As a result, we obtained CCI events for each dataset across the whole-cell, cytoplasm, and nucleus. We used Benjamini-Hochberg method to adjust the *P* -values. An adjusted P-value threshold of ≤ 0.05 in atleast one of the compartments was used as a permissive primary filter to avoid prematurely excluding low-abundance subcellular signaling events. We then used the whole-cell communication scores across these nine datasets for each CCI event as model inputs along with the other features detailed in “Feature scaling and encoding section”. CCIDeconv assigns a CCI event to class ***x*** (separable class), if there are any significant communication scores in the cytoplasm or nucleus. Otherwise, it assigns them to class ***o*** (terminal class). We used these class labels as ground truth for classification, whereas the cytoplasm and nucleus communication scores were used as ground truth to train the regression models.

### Evaluation of candidate models for CCIDeconv

#### Leave one group out cross-validation

We evaluated CCIDeconv using leave one group out cross-validation (LOGO-CV) (Pedregosa et al., 2011) on the nine sST datasets. We used the SP procedure to assess CCIDeconv performance on the sST dataset and the ScP procedure to assess the applicability of CCIDeconv to single-cell datasets. We conducted scaling and target encoding steps separately for the training and testing data to prevent information leakage. Hyperparameter tuning was done only in the training folds in each cross-validation.

#### Model selection from different baselearner configurations and encoding location features

We evaluated seven different hierarchical models with different base learning configurations and feature encoding strategies of the subcellular location features from CellChatDB and HPA. The details of the various model specifications are given in Supplementary Table 1. We selected the best and balanced model using LOGO-CV.

**Table 1.** Evaluation of CCIDeconv (LOGO-CV) across the nine sST datasets using SP procedure.

| Metric | Mean | Median | Std Dev | Std Err | IQR |
|---|---|---|---|---|---|
| AUC | 0.79 | 0.80 | 0.11 | 0.04 | 0.12 |
| Recall (macro) | 0.69 | 0.67 | 0.09 | 0.03 | 0.09 |
| *R*<sup>2</sup> cytoplasm | 0.87 | 0.88 | 0.07 | 0.02 | 0.05 |
| *R*<sup>2</sup> nucleus | 0.8 | 0.80 | 0.14 | 0.04 | 0.13 |
| NRMSE cytoplasm | 0.34 | 0.35 | 0.12 | 0.04 | 0.08 |
| NRMSE nucleus | 0.41 | 0.44 | 0.17 | 0.06 | 0.16 |
| **Composite metric** | **0.73** | **0.75** | **0.06** | **0.02** | **0.1** |

#### Evaluation metrics

For each stage of the hierarchical framework, we evaluated model performance independently using unknown test data.

##### Classification

We calculated accuracy, AUC, and Macro recall across all input CCI events to evaluate the assignment of terminal (class ***o***) and separable (class ***x***) labels.

##### Regression

For Class ***x*** events, we assessed the deconvolution precision by measuring *R*^2^ and Normalised Root Mean Squared Error (NRMSE) of predicted *versus* ground-truth scores for the nucleus and cytoplasm. We calculated NRMSE by normalising the mean squared error by the standard deviation of the response, i.e., communication scores in nucleus or cytoplasm.

##### Composite Assessment

We used a composite metric, calculated as the geometric mean of all classification and regression metrics, for a holistic assessment of the total framework performance. We used the geometric mean to ensure that poor performance in any single component penalises the overall score, preventing a high score in classification or regression from masking weakness in the other.

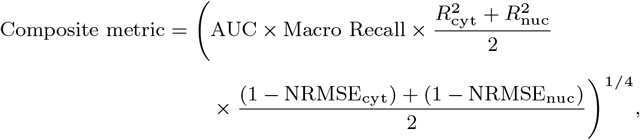

where NRMSE_cyt_, NRMSE_nuc_, 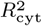, and 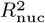 represent the NRMSE and *R*^2^ values of cytoplasmic and nuclear regression respectively.

#### Robustness analysis

To explore the robustness of CCIDeconv with respect to the number of datasets, we trained on every possible combination of eight datasets with the ninth dataset set aside as the test data. This resulted in 255 training combinations for each dataset. We conducted scaling and target encoding steps within each combination of datasets to prevent information leakage. We used both SP and ScP procedures for each combination and assessed performance using the composite metric.

### Application to scRNA-seq data

We applied CCIDeconv to a lung cancer single-cell dataset (Supplementary Methods 1.2.3.) by training on seven sST datasets using the ScP procedure. We removed the two remaining sST datasets (Brain (AD) and Lung) because their compartment-specific communication was primarily driven by the spatial features. We deconvoluted the whole-cell CCI to obtain cytoplasm and nucleus specific CCI predictions. We performed a direct comparison of cytoplasm and nucleus CCI by calculating their *log*_2_ difference.

### Independent validation and compartment-specificity analysis

We compiled independent localisation evidence from reviewed UniProtKB/Swiss-Prot annotations, non-HPA and CellChatDB COMPARTMENTS records and Reactome. We excluded HPA-derived annotations and CellChatDB to reduce circularity. Evidence comprised direct protein-localisation annotations, and indirect evidence supporting localisation through mechanistic or pathway associations. We quantified the overlap between the top 10 and top 20 events from the SP and ScP procedures and scRNA-seq predictions against these independent resources.

To assess whether our model identifies compartment-specific events, we compared the top-ranked nuclear and cytoplasmic events from the SP and ScP procedures. We quantified the proportion of events with independent nuclear-localisation support in each group and calculated the difference in proportions. We assessed statistical significance using a one-sided, dataset-stratified permutation test with 10,000 iterations. We held events shared between the ranked groups fixed and restricted permutations to discordant events within each dataset. We calculated P values using a plus-one correction and estimated confidence intervals for the difference in proportions and fold enrichment using paired bootstrap resampling.

### Benchmarking performance of CCIDeconv

We compared CCIDeconv with three baseline allocation models and flat classification and regression only models to assess whether subcellular communication scores could be estimated using simpler allocation strategies

- **Direct assignment:** The whole-cell communication score was assigned directly to both compartments without deconvolution:

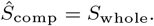

where *S*_whole_ is the whole-cell communication score and *Ŝ*_comp_ is the predicted/ allocated compartment-specific communication score

- **HPA proportional split:** The whole-cell score was divided between the cytoplasm and nucleus using compartment-specific localisation weights derived from the Human Protein Atlas (HPA):

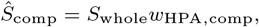

where *w*_HPA,comp_ is the HPA-derived localisation weight for the corresponding compartment, with the cytoplasmic and nuclear weights summing to one. Each compartment therefore receives a proportion of the whole-cell score determined by its localisation weight.

- **Mean allocation:** The mean ratio of compartment-specific to whole-cell scores was estimated from the training datasets and applied to the held-out dataset:

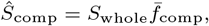

where 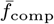 is the mean compartment-to-whole-cell score ratio estimated from the training data. This fraction was applied to the held-out whole-cell score to estimate the corresponding compartment score.

We also benchmarked CCIDeconv by comparing it against flat four-class classification or independent compartment regressions alone. We evaluated the flat-baseline models multiclass classification using One-vs-Rest (OvR) macro-averaged ROC Area Under the Curve (AUC) and macro-averaged recall, which compute individual class performance metrics against the remaining classes before taking their unweighted average. All methods were evaluated across nine leave-one-dataset-out folds by comparing allocated/ predicted compartment scores with the corresponding ground-truth sCCI scores.

## Results

### Interactions between ligands and receptors vary between the subcellular compartments

We hypothesised that CCI varies across subcellular compartments due to subcellular expression differences (Fig. 1A). To test our hypothesis, we computed communication scores in cytoplasm and nucleus across nine sST datasets covering Brain (AD and GL datasets), Lung (2 datasets), Lymph Node, Ovarian, Pancreas, Skin, and Breast Cancer tissues (Supplementary methods 1.2.1.). We observed marked differences in both the number of communicating CCI events, hereafter referred to as LR pairs (Fig. 1B, Supplementary Fig. 2), and their communication scores in all datasets (Fig. 1C, Supplementary Fig. 3 and 4). We identified 3,273 significant communicating LR pairs across the nine datasets. These LR pairs recur, but are unique when defined based on the interacting cell-types. These LR pairs were significant in at least one of the whole-cell, cytoplasm or nucleus compartments (BH-adjusted *P* -value ≤ 0.05; Supplementary Table 2). We identified 168 common pairs between whole-cell and nucleus-nucleus communication and 568 common pairs between whole-cell and cytoplasm-cytoplasm communication, indicating differences in sCCI. We also observed differences in communication score between cross-compartment sCCI (cytoplasm to nucleus and nucleus to cytoplasm; Supplementary Fig. 4).

**Figure 2.**
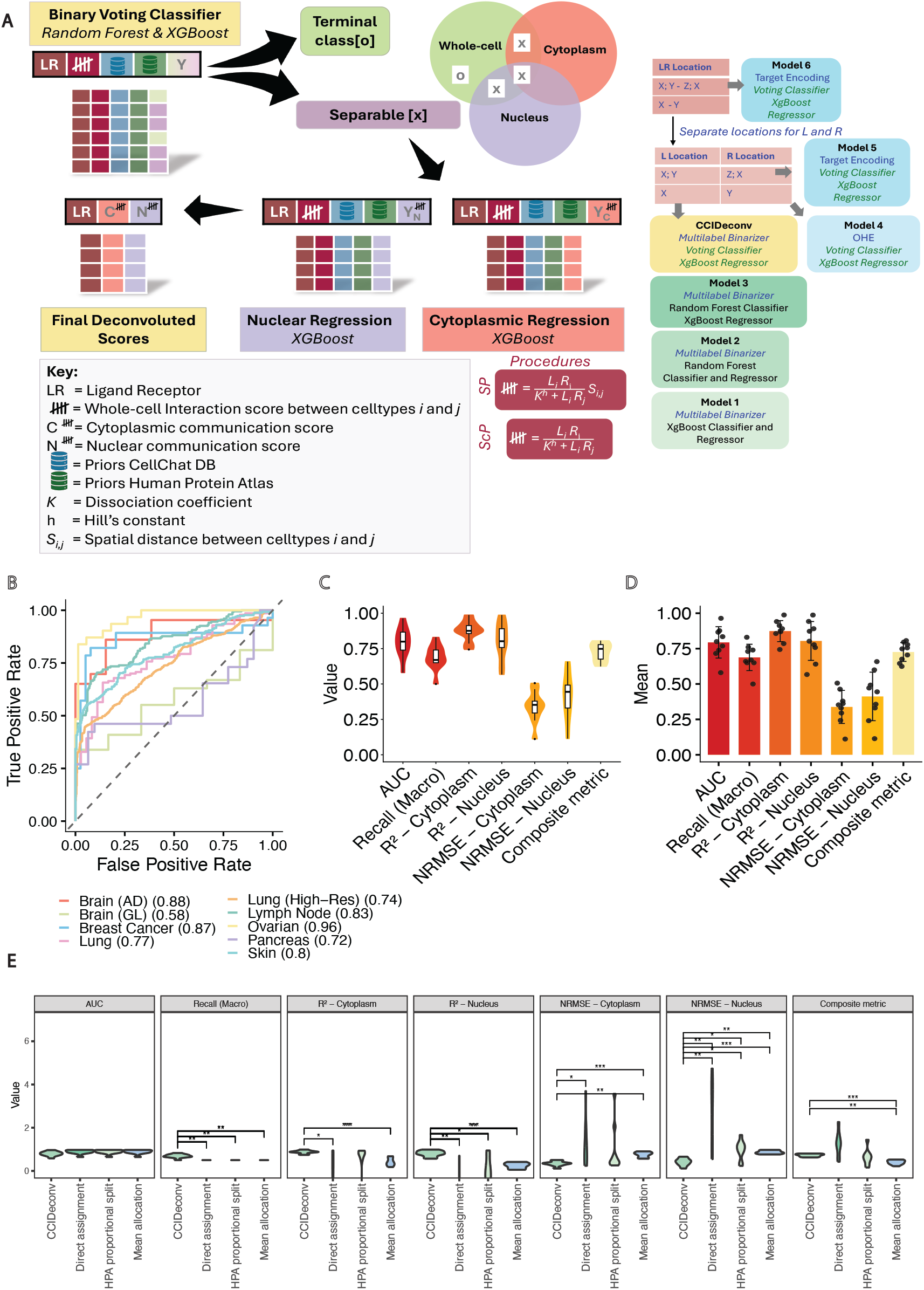
Training and evaluation of CCIDeconv, a hierarchical model for predicting subcellular communication scores. A. Schematic of the hierarchical model architecture of CCIDeconv, combining a classifier and a regressor to predict communication scores in subcellular compartments. B. ROC curves showing the classification performance of the model on nine datasets, with the area under the curve (AUC) for each dataset indicated in the legend. C. Violin plots showing the distribution of performance metrics. D. Barplots representing the mean of performance metrics across datasets with standard error bars. E. Benchmarking the performance of CCIDeconv under the SP procedures with the different baseline comparisons. Statistical comparisons between CCIDeconv and each baseline were performed using paired Wilcoxon signed-rank tests, with P-values adjusted for multiple comparisons using the Benjamini–Hochberg (BH) method. Statistically significant differences are indicated in the figure.

**Figure 3.**
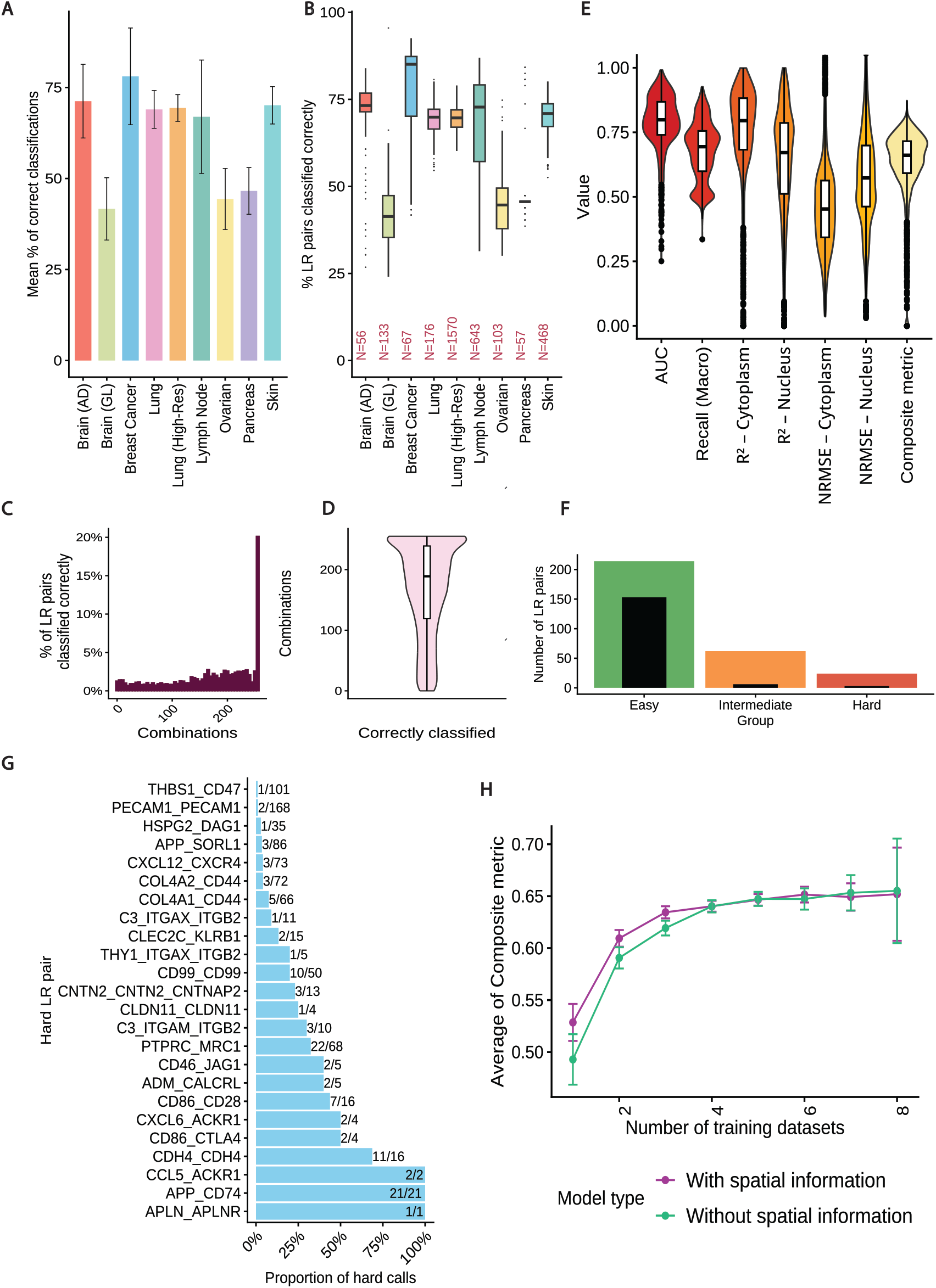
Assessing model stability by training each dataset with varying combinations of the remaining datasets. A, B. Mean percentage (A) and distribution (B) of significant ligand–receptor (LR) pairs correctly classified when each dataset was trained with 255 different combinations of the other eight datasets. C, D. Number of training combinations in which each significant LR pair across the nine datasets was correctly classified: (C) histogram of correctly classified LR pairs; (D) violin plots showing the distribution of the number of combinations in which each LR pair was correctly classified. E. Violin plot of performance metrics across the 255 training combinations for all nine datasets (outliers with cytoplasmic and nuclear *R*^2^ *≤* 0 are not shown). F. Counts of LR pairs consistently assigned to diffculty groups. Bars represent the number of significant LR pairs that were classified as Easy (≥75%), Intermediate (25–75%), or Hard (*≤* 25%) based on the proportion of training combinations in which they were correctly classified across all datasets. Gray overlays indicate LR pairs that were always assigned to the same difficulty group across all combinations. G. Bar plot of LR pairs classified as Hard; fractions shown to the right of each bar indicate the proportion of combinations in which each LR pair was labeled as Hard. H. Composite metrics for models trained with SP and ScP procedure across different dataset combinations. A total of 4,590 models were trained (each of the nine datasets trained across 255 combinations of other datasets with both the SP and ScP procedures); plot shows mean composite metrics with error bars.

**Figure 4.**
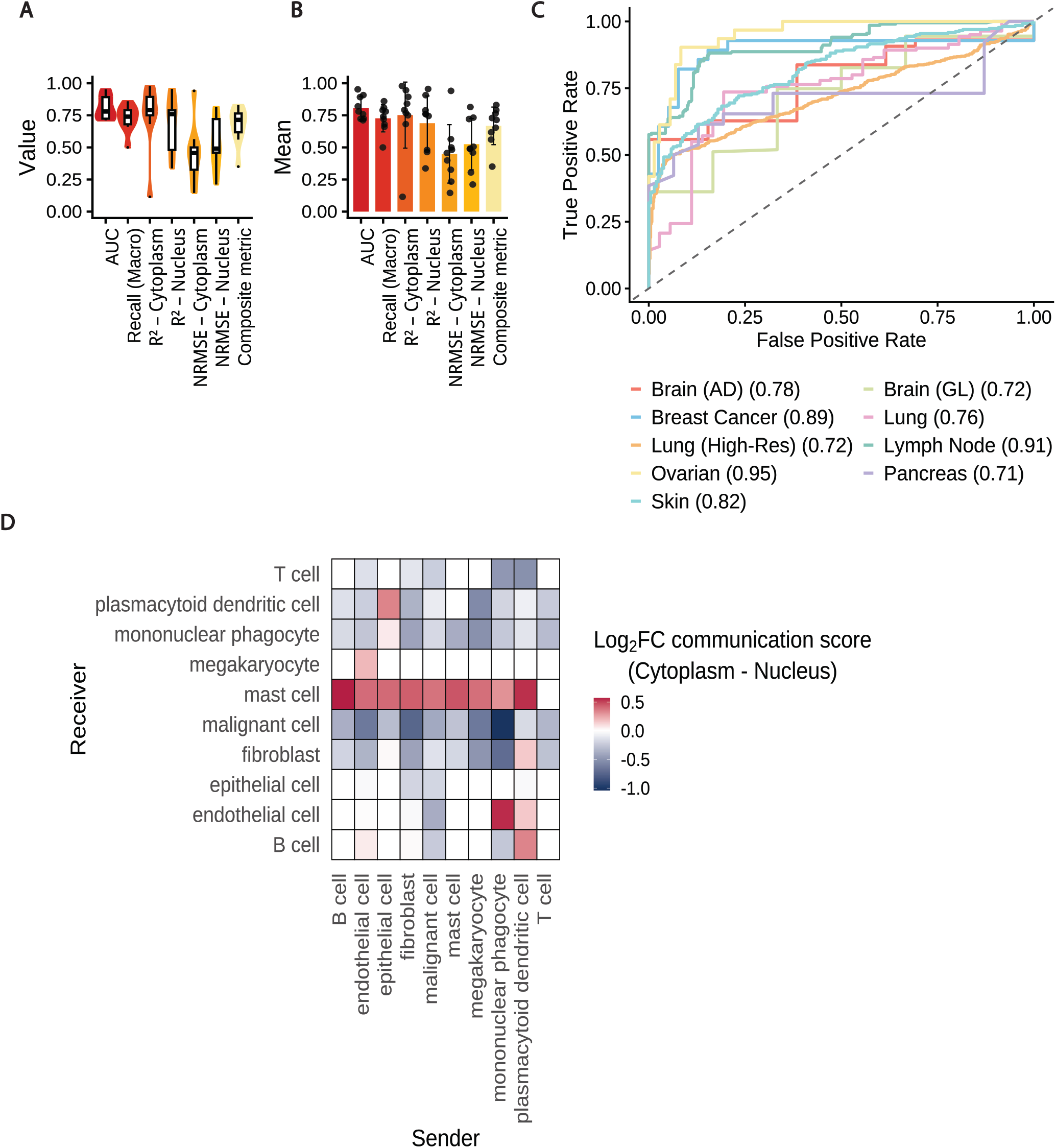
Training the CCIDeconv through ScP procedure to test applicability to single-cell data. A - C Performance metrics on training the model without the spatial features: (A) Violin plots representing the distribution of the metrics across the datasets (B) Bar plots representing the mean across datasets with standard error bars and (C) ROC curves showing the classification performance of the model on nine datasets, with the area under the curve (AUC) for each dataset indicated in the legend. D. Heatmap showing the difference in the communication scores in the different subcellular compartments when the model is applied on a single-cell data.

As a biological consistency check, (*GJA1* -*GJA1*) communication was detected predominantly in the cytoplasmic compartment in the Brain (AD) dataset, consistent with the localisation of the gap junction protein *GJA1*. Several nucleus-biased events, including those involving *PTPRC* and *KLRB1*, were also supported by HPA nucleoplasmic annotations. More broadly, highly ranked compartment-associated CCI events showed concordance with established protein localisation annotations (Supplementary Table 3).

We further assessed if different components of CellChat communication score capture distinct, compartment-specific patterns of CCI. We performed Principal Component Analysis (PCA) on various components of the CellChat communication score (i.e., Hill function, Spatial, Agonist, and Antagonist components), which showed that LR pairs separated based on the compartment where they were detected (Fig. 1D). We found compartment-specific communication patterns were mostly driven by the Hill function of LR expression (Supplementary Fig. 5). Together with the observed differences in the number of communicating LR pairs, their communication scores, and PCA-based separation, these results indicate that CCI varies between cytoplasm and nucleus.

### CCIDeconv’s hierarchical modeling enables robust subcellular deconvolution of spatial communication scores

Having established that LR communication scores differ between subcellular compartments, we next evaluated whether CCIDeconv could deconvolute whole-cell communication scores into nuclear- and cytoplasmic-associated components (Fig. 2A). Using LOGO-CV with the SP procedure, CCIDeconv achieved a median composite metric of 0.75 across the nine datasets (Fig. 2B-D; Table 1). Classification performance was moderate to strong, with a mean AUC of 0.79 and macro recall of 0.69. Regression performance was higher for cytoplasmic-associated scores (*R*^2^ = 0.87, NRMSE = 0.34) than for nuclear-associated scores (*R*^2^ = 0.80, NRMSE = 0.41; Supplementary Fig. 6 and 7). To evaluate our performance further, we benchmarked our hierarchical model with different baseline comparisons and flat baseline models (Fig. 2E, Supplementary Fig. 8 and 9) and found CCIDeconv outperformed all the comparisons, while achieving similar performance to an end-to-end regression approach evaluated across LOGO-CV, supporting the value of the hierarchical framework (Supplementary Fig. 9). We also performed a sensitivity analysis using 3,193 events significant in the whole-cell alone at adjusted P value ≤0.05, evaluated using LOGO-CV with the SP procedure. The median composite metric was 0.71, compared with 0.75 in the primary analysis (Supplementary Fig. 10).

To validate compartment specificity, we identified independent localisation support for 169/180 (93.8%) cytoplasmic and 133/180 (73.9%) nuclear predictions among the top 20 events across all nine datasets (Supplementary Fig. 11) in the SP procedure. We further assessed compartment specificity across our predictions and found that top-ranked nuclear events had significantly greater nuclear-localisation support than top-ranked cytoplasmic events for both the top 10 (*P* = 0.0002) and top 20 predictions (*P* = 0.0201; Supplementary Table 4). To assess whether location features drove the improved performance, we performed ablation analysis, which showed that performance declined most when whole-cell communication scores were removed, indicating that localisation features contributed to, but did not solely drive, model performance (Supplementary Table 5). Training and prediction required less than 10 minutes per held-out dataset on a standard laptop. Together, these results show that CCIDeconv can estimate compartment-associated communication scores in unseen datasets, although performance was consistently stronger for cytoplasmic than nuclear predictions.

### CCIDeconv exhibits consistent performance across diverse tissue types

We performed robustness analysis on CCIDeconv using the SP procedure to assess its stability across different tissue types (Materials and methods; Robustness analysis). We observed that a median of 67.3% of LR pairs were classified correctly across all training combinations and datasets (IQR: 45.6%–73.3%; Fig. 3A–D), with a median AUC of 0.80 (Fig. 3E). Approximately 20% of the LR pairs were always correctly classified, and less than 5% were incorrectly classified, across all training combinations (Fig. 3C). Most of the incorrect classifications belonged to the Brain (GL), Ovarian, and Pancreas datasets (Supplementary Fig. 12A and B), which also had a lower proportion of correctly classified LR pairs (Fig. 3A).

To further examine the LR pair classifications, we labeled LR pairs as “Easy” if they were correctly classified in at least 75% of the combinations, “Hard” if they were classified correctly in less than 25% of the combinations, and “Intermediate” otherwise. We found 214 Easy, 24 Hard, and 62 Intermediate LR pairs (Fig. 3F). Out of these, 153 LR pairs were classified correctly across all training combinations and only 3 were always classified incorrectly. For example, the APP CD74 LR pair was detected 21 times irrespective of cell and tissue type and was always classified incorrectly (Fig. 3G).

Analysis of the regression component of CCIDeconv showed that the cytoplasmic *R*^2^ was high (median= 0.80) with a low NRMSE (median = 0.45) (Fig. 3E, Supplementary Table 6). Although the NRMSE-nucleus was higher than expected (median = 0.57), the *R*^2^-nucleus was considerably higher (median = 0.67) indicating that CCIDeconv achieves good regression performance across different tissue types. This was further supported by a median composite value (0.66) and consistent performance metrics across all datasets (Supplementary Fig. 12 C and D). To investigate factors underlying the slightly weaker nuclear predictions, we correlated nuclear regression metrics with six dataset characteristics capturing nuclear mRNA sparsity (Nuclear detected genes, Nuclear zero-count fraction), nuclear–cytoplasmic expression distributions (Nuclear detection ratio, N/C expression ratio), and technical dropout rates (Nuclear–cytoplasmic dropout gap) across the LOGO-CV and robustness settings of both the SP and ScP procedures. The nuclear–cytoplasmic dropout gap showed positive associations with both performance differences (Spearman (*ρ*) up to 0.73). The N/C expression ratio showed moderate negative associations with several metrics, whereas the other characteristics showed weaker or less consistent associations. This shows that nuclear prediction accuracy may depend on differential transcript detection between compartments (Supplementary Fig. 13). However, the top-ranked nuclear and cytoplasmic predictions retain independent biological support (Supplementary Fig. 11). Therefore, we conclude CCIDeconv maintained broadly stable performance across datasets.

### Spatial information improves CCIDeconv performance in low training data settings

We performed robustness analysis using both the SP and ScP procedures (Materials and methods; Robustness analysis) to determine the importance of spatial features in deconvoluting the communication score. We observed that the composite metric was lower in models trained without the spatial features when the number of training datasets was less than 4 (Fig. 3H). However, as the number of training datasets increased, models trained with or without spatial features achieved similar results. A similar pattern was also observed for AUC and *R*^2^ in cytoplasm (Supplementary Fig. 14). In contrast, macro recall and *R*^2^ in nucleus showed the opposite pattern. Nevertheless, NRMSE in both compartments was high in models trained without the spatial features when there were fewer training datasets, but dropped as the number of training datasets increased. Therefore, when training with a higher number of datasets, spatial information does not improve performance, indicating CCIDeconv could be applied to datasets with no spatial information.

### Application of CCIDeconv to non-spatial single-cell data identifies communication patterns consistent with known biology

To assess the applicability of CCIDeconv to non-spatial single-cell datasets, we trained the model using LOGO-CV and the ScP procedure. We observed a median composite metric of 0.71 during the training (Fig. 4A-C, Supplementary Table 7). We identified independent support for the localisation evidence in external databases for 168/180 (93.3%) top 20 and 129/180 (71.7%) top 20 cytoplasmic and nuclear predictions across all nine datasets respectively for the ScP procedure (Supplementary Fig. 11). Top-ranked nuclear events had significantly greater independent nuclear-localisation support than top-ranked cytoplasmic events (P = 0.0001 for both the top 10 and top 20 nuclear predictions; Supplementary Table 4). To further investigate the platform transferability, we tested CCIDeconv SP with a CosMx lung cancer dataset (Supplementary method 1.2.3.). Although the classification performance was not robust (AUC = 0.53, macro recall = 0.45, cytoplasmic R^2^ = 0.61, nuclear R^2^ = 0.48), the regression results remained informative. Furthermore, domain-shift stress testing (Supplementary Fig. 15) showed that both regression and classification were robust to most distributional changes, with classification primarily affected by mean shifts in whole-cell communication scores, supporting the potential application of CCIDeconv to scRNA-seq data. Therefore, we tested the ScP procedure of CCIDeconv on a scRNA-seq lung cancer dataset (Supplementary Methods 1.2.4.) to assess whether our model could recover known communication patterns from a non-spatial dataset. We also found that among the nine sST training datasets the Lung Cancer and Brain (AD) datasets showed poor correlation between the compartment-specific communication scores and the whole-cell communication score (*r ≤* 0.7), indicating that spatial information was altering the correlation (Supplementary Fig. 16 A and B). Therefore, we used the remaining seven datasets as training datasets to predict communication patterns in the scRNA-seq lung cancer dataset. We observed a high communication score between fibroblasts and mast cells in the cytoplasm indicating potential communication between these two cell lineages (Fig. 4D, Supplementary Fig. 16C), which has been previously reported to occur through cell contact (Larsson-Callerfelt et al., 2021). Our model identified putative nuclear-biased communication scores between mononuclear phagocytes and malignant cells, likely driven by the *FN1* –*CD44* signaling event. Functional CD44 protein translocates to the nucleus via endocytic pathways (Rios de la Rosa et al., 2019). We also observed that CD44 participates in nuclear transcriptional regulation according to information in the Reactome Pathway Knowledgebase (R-HSA-1031716, Ragueneau et al. (2026)), further supporting our prediction. CCIDeconv was able to capture this axis at the transcript level by isolating nuclear mRNA enrichment. We identified independent support for the localisation evidence in external databases for all (100%) top 20 and 19/20 (95%) top 20 cytoplasmic and nuclear predictions respectively for this lung cancer scRNA seq dataset. Together, these examples support the potential application of CCIDeconv for inferring candidate compartment-biased communication patterns from non-spatial scRNA-seq datasets.

## Discussion

In this study, we demonstrate that CCI is not uniformly distributed across subcellular compartments and that incorporating subcellular information can provide additional resolution beyond conventional cell-level CCI analyses. By leveraging subcellular spatial transcriptomics data, CCIDeconv deconvolutes whole-cell communication scores into compartment-specific communication and demonstrates the potential for applying these predictions to non-spatial single-cell transcriptomic data. CCIDeconv included HPA-validated localisation features to capture biologically meaningful signals. Across nine independent sST datasets, CCIDeconv showed consistent performance, suggesting that subcellular communication patterns are suffciently reproducible to support transfer learning across tissues. To the best of our knowledge, this is the first method to deconvolute whole-cell level communication into subcellular CCI, leaving no existing methods for direct comparison.

Despite the use of CellChatDB to characterise LR interactions, because of its broad coverage of curated signaling pathways, CCIDeconv is not restricted to a specific interaction database. Alternative resources such as OmniPath (Türei et al., 2016) or CellPhoneDB (Troulé et al., 2025) could be readily incorporated into the framework. This flexibility allows CCIDeconv to adapt as ligand-receptor knowledge bases continue to expand and improve.

Here we have focused on the task of deconvoluting CCI to the nucleus or the cytoplasm. Cells contain many different subcellular compartments of interest including the mitochondria, golgi body, or plasma membrane. While it is desirable to develop an analytical approach that can predict the origin of CCI to all subcellular compartments, many existing sST datasets are annotated to the resolution of the nucleus and cytoplasm only. One potential approach to ameliorate the lack of annotated subcellular reference data is to use the local gene expression abundances themselves as evidence for presence of cellular substructures, for example, subcellular compartments enriched in mitochondrial genes may truly represent the mitochondria of the cell, and be annotated as such. This is an exciting avenue that could be explored in future work.

Across the nine datasets studied, we did not identify significant LR pairs uniquely associated with cross-compartment communication between the nucleus and cytoplasm. One possible explanation is the limited gene coverage of current 10X Xenium panels (approximately 5,000 genes), relative to whole-transcriptome measurements, which might have reduced the ability to detect low-abundance signalling molecules involved in compartment-specific communication. In particular, key signalling genes, including agonists and antagonists, may not be suffciently represented or expressed to enable reliable detection of cross-compartment interactions. Future studies using broader transcriptomic panels and improved subcellular annotations may provide greater power to identify such signalling relationships.

Ideally, CCI should be evaluated using proteomics data because proteins directly mediate cellular function. However, subcellular spatial proteomics remains sparse and has limited coverage, and most current methods, including CCIDeconv, rely on transcriptomics as a proxy for assessing CCI. As a result, the inferred communication scores reflect transcript-level association rather than direct protein binding or signaling, and therefore, should be interpreted cautiously. Nevertheless, CCIDeconv can prioritize LR pairs and their likely subcellular context, providing candidates for follow-up validation using spatial proteomics or other experimental approaches.

In conclusion, our findings highlight that CCI varies between subcellular compartments. We developed a hierarchical model, CCIDeconv, to attribute the whole-cell communication scores into their cytoplasm and nucleus scores. The predictions generated by CCIDeconv provide a framework for exploring the subcellular context of CCI and generating hypotheses regarding disease-associated signalling pathways.

## Supporting information

Supplementary Methods and Results

## Key Points

- We observed differences in CCI between subcellular compartments, however current CCI analysis methods assess cell communication at the whole-cell-level, without considering the sub-cellular localisation of ligands and receptors.
- CCIDeconv is a hierarchical machine learning framework that deconvolutes whole-cell communication to assign each compartment (nucleus or cytoplasm) subcellular communication scores.
- CCIDeconv enhances biological fidelity by including validated protein localisation information from the Human Protein Atlas (HPA) and CellChatDB as prior categorical features during deconvolution.
- CCIDeconv achieves robust generalisability across nine publicly available human sST datasets via leave-one-dataset-out cross-validation, yielding a median composite metric of 0.75 and strong mean *R*^2^ values of 0.87 (cytoplasmic) and 0.80 (nuclear).
- CCIDeconv can be trained using two flexible procedures, a Spatial procedure (SP) trained with the spatial distance features and a Single-Cell Procedure (ScP) designed for non-spatial data. We demonstrate that spatial features are especially valuable when training data is limited, however, as the number of training datasets increases, the Single-Cell Procedure (ScP) achieves near similar performance and this provides a potential approach for application to non-spatial single-cell transcriptomic data.

## Acknowledgments

We thank our colleagues in the University of Sydney School of Mathematics and Statistics, Charles Perkins Centre, and Sydney Precision Data Science Centre for their support and intellectual engagement.

## Author contributions statement

S.G. and J.Y. conceived the study. R.J. developed the method, performed data analysis, and wrote the initial draft of the manuscript under the supervision of J.Y., S.G., and P.P. All authors read and approved the final version of the manuscript.

## Data and code availability

No new data were generated in this study. All analyses were performed using publicly-available datasets, as described in the Supplementary Methods section, with links and references provided there. Model weights can be downloaded from https://doi.org/10.5281/zenodo.24037096.

Analysis and figure generation were performed using R (version 4.5.1) and Python (version 3.13.1). The model and scripts are available at https://github.com/SydneyBioX/CCIDeconv.

## Biographical note

The authors of CCIDeconv are bioinformaticians from the Sydney Precision Data Science Centre at the University of Sydney. CCIDeconv provides researchers the opportunity to study subcellular communication.

