## Supplementary Methods and Results for "CCIDeconv: Hierarchical model for deconvolution of subcellular cell-cell interactions in single-cell data"

#### Authors and Affiliations

Rojashree Jayakumar[1,2,3], Pratibha Panwar[1,2, 3], Jean Yee Hwa Yang[1,2,3], Shila Ghazanfar[1,2,3]

[1]School of Mathematics and Statistics, The University of Sydney, Camperdown, New South Wales, 2006, Australia.

[2]Sydney Precision Data Science Centre, The University of Sydney, Camperdown, New South Wales, 2006, Australia.

[3]Charles Perkins Centre, The University of Sydney, Camperdown, New South Wales, 2006, Australia.

### 1. Supplementary methods

#### 1.1. Hyperparameter tuning

For the Random Forest classifier, optimization was performed over the *max_samples* (0.1–1), *max_features* (0.3–1), *n_estimators* (150–300), *max_depth* (5–30), *min_samples_split* (2–10) and *min_samples_leaf* (2–10).

Similarly for the XGBoost classifier, optimization was over the *max_depth* (6–10), *learning_rate* (0.01–0.3), *n_estimators* (100–300), *reg_alpha* (0-1), *colsample_bytree* (0.5-1), *scale_pos_weight* (1,100) and *gamma* (0–1). Model performance during optimisation for the classifiers was assessed using cross-validated F1-score.

For the Random Forest regressor, the following hyperparameters were optimised; *max_depth* (5-30), *n_estimators* (100-300), *max_samples* (0.5-1), *min_samples_split* (2-20) and *min_samples_leaf* (1-10).

For the XGBoost regressor, the following hyperparameters were optimized, *max_depth* (10-20), *learning_rate* (0.01-0.1), *n_estimators* (100-300), *reg_alpha* (0-1), *min_child_weight* (1-3), *colsample_bytree*(0.8-1) and *gamma* (0-3). Model performance during optimisation for the regressors was assessed using cross-validated $R^2$.

For all the models, the optimisation was initialised with three random evaluations and subsequently run for 30 iterations. The hyperparameters corresponded to the parameter set that maximised the optimisation objective. The objective function of the XGBoost classifier was *binary:logistic* and the regressor was *reg:squarederror.*

#### 1.2. Datasets

##### 1.2.1. 10X Xenium Datasets

We processed nine publicly available subcellular spatially resolved transcriptomics datasets. All the datasets were accessed from 10X genomics. We chose datasets matching at least one of the following criteria:

1. High transcript density - Median number of transcripts per cell > 100
2. High-plex profiling - Utilisation of Xenium 5K panel
3. Biological diversity - Datasets representative of diverse tissue types

**[i] Brain datasets:** We had two adult human brain datasets from an Alzheimer's disease patient (Brain -AD) and glioblastoma (Brain -GL) patient. Both the datasets were FFPE-preserved tissue blocks. The Alzheimer's dataset has 44,955 cells and 216 median transcripts per cell. The glioblastoma dataset has 40,887 cells and 216 median transcripts per cell. The panel used was Xenium Human Brain Gene Expression Panel along with an additional 65 genes. The data was downloaded from https://www.10xgenomics.com/datasets/xenium-human-brain-preview-data-1-standard on April 2025.

**[ii] Breast dataset:** The raw expression data from FFPE-preserved tissue blocks of human breast cancer has a total of 699,110 cells and 51 median transcripts per cell. The Xenium Prime 5K Human Pan Tissue and Pathways Panel was used along with 100 custom genes. The data was downloaded from https://www.10xgenomics.com/datasets/xenium-prime-ffpe-human-breast-cancer on April 2025.

**[iii] Lymph node dataset:** The raw expression data from FFPE-preserved tissue blocks of human reactive lymph nodes has a total of 708,983 cells and 255 median transcripts per cell. The Xenium Prime 5K Human Pan Tissue and Pathways Panel was used along with an additional 377 genes. The data was downloaded from https://www.10xgenomics.com/datasets/preview-data-xenium-prime-gene-expression on April 2025.

**[iv] Lung datasets:** Two lung cancer datasets were used in our analysis.
(i) Lung: The raw expression data from FFPE-preserved tissue blocks of adult human Lung Adenocarcinoma tissue has a total of 162,254 cells and 46 median transcripts per cell. The Xenium Human Multi-Tissue and Cancer Panel (377 genes) was used. The data was downloaded from https://www.10xgenomics.com/datasets /preview-data-ffpe-human-lung-cancer-with-xenium-multimodal-cell-segmentation-1-standard on April 2025.

(ii) Lung (High res): The raw expression data from FFPE-preserved tissue blocks of adult human Lung Adenocarcinoma tissue has a total of 278,328 cells and 242 median transcripts per cell. The Xenium Human Lung Gene Expression Panel and Xenium Prime 5K Human Pan Tissue and Pathways Panel was used. This dataset is labelled as Lung (high-res). The data was downloaded from https://www.10xgenomics.com/datase ts/xenium-human-lung-cancer-post-xenium-technote on April 2025.

**[v] Ovarian dataset:** The raw expression data from fresh frozen tissue of human ovarian adenocarcinoma has a total of 1,157,659 cells and 1,401 median transcripts per cell. The Xenium Prime 5K Human Pan Tissue and Pathways Panel was used. The data was downloaded from https://www.10xgenomics.com/datasets/xenium-prime-fresh-frozen-human-ovary on April 2025.

**[vi] Pancreas dataset:**The raw expression data from FFPE-preserved tissue blocks of human pancreatic cancer tissue has a total of 190,765 cells and 114 median transcripts per cell. TheXenium Human Multi-Tissue and Cancer Panel (377 genes) was used. The data was downloaded from https://www.10xgenomics.com/da tasets/pancreatic-cancer-with-xenium-human-multi-tissue-and-cancer-panel-1-standard on April 2025.

**[vii] Skin dataset:** The raw expression data from FFPE-preserved tissue blocks of human reactive lymph node has a total of 112,551 cells and 306 median transcripts per cell. The Xenium Prime 5K Human Pan Tissue and Pathways Panel besides the following genes *ADGRG1, NMBR, OSMR, OSTF1, and RGS8* were used. The data was downloaded from https://www.10xgenomics.com/datasets/xenium-prime-ffpe-human-skin on April 2025.

###### 1.2.2. Reference datasets

**[i] Cancer reference dataset:** To annotate the cancer cells, single cell cancer atlas dataset containing eight different human cancer tissues (lung, breast, melanoma, liver, uvea, colorectum, skin epidermis and ovaries) from the study by Guimarães et al. (2024) was used. The integrated dataset was downloaded from https://cellxgene.cziscience.com/collections/3f7c572c-cd73-4b51-a313-207c7f20f188 on April 2025.

**[ii] Glioblastoma reference dataset:** To annotate the glioblastoma disease dataset, from the study by Ruiz-Moreno et al. (2022) was used. The full dataset was downloaded from https://cellxgene.cziscience.com /collections/999f2a15-3d7e-440b-96ae-2c806799c08c on April 2025.

**[iii] Alzheimer's reference dataset:** To annotate the Alzheimer's disease dataset, from the study by Pan et al. (2024) was used. The full dataset was downloaded from https://cellxgene.cziscience.com/collections/7 c4552fd-8a6d-4da3-9854-2dfa8baca8bf on April 2025.

**[iv] Lymph Node reference dataset:** To annotate the lymph node dataset, Single-cell RNA-seq data were obtained from the Human Expression (HE) organs reference dataset (2020 release), accessed through the scRNAseq package (Risso and Cole, 2026) using the fetchDataset interface (dataset version 2023-12-21 and path lymph_node) on April 2025.

**[v] Pancreas reference dataset:** To annotate pancreatic cell types, the pancreasref.SeuratData Azimuth reference (Hao et al., 2024) was used. This reference consists of a curated single-cell RNA-seq Seurat object with annotated human pancreatic cell types and was accessed through the SeuratData framework (version 1.0.0) on April 2025.

##### 1.2.3. CosMx Dataset

The Lung 13 dataset was downloaded from the Bruker Spatial Biology website (https://brukerspatialbiology.com/resources/smi-ffpe-dataset-lung13-data/) on October 2026.

##### 1.2.4. scRNA-seq Dataset

scRNA-seq data for human lung tissue was obtained from the Zilionis et al. (2019) study. The dataset was accessed through the scRNAseq package (Risso and Cole, 2026) using the fetchDataset interface (dataset version 2023-12-20 and path human) on April 2025.

#### 1.3. Feature-ablation analysis

We performed a controlled feature-ablation analysis on both the SP and ScP procedures of CCIDeconv to assess the contribution of subcellular localisation annotations, whole-cell communication score and LR identity to predictions. Seven feature configurations were evaluated: (i) the full model, (ii) removal of CellChatDB localization features, (iii) removal of HPA localization features, (iv) removal of all localization features, (v) removal of the ligand–receptor pair identifier, (vi) removal of the whole-cell communication score and (vii) removal of the molecular information while retaining source and target cell types and the remaining ligand/receptor and spatial features. The hierarchical classifier comprised a calibrated Random Forest (200 trees, maximum depth 15, minimum samples split of 5, minimum samples leaf of 3, maximum features of 0.5, and maximum samples of 0.8) and XGBoost classifier (200 trees, maximum depth 8, learning rate 0.1, minimum child weight 5, gamma 1.0, column subsampling of 0.8, and L1 regularization of 0.5). Cytoplasmic and nuclear regressors used 150 trees, maximum depth 12, learning rate 0.05, minimum child weight 2, gamma 1.0, column subsampling of 0.9, and L1 regularization of 0.5. These calibrated hyperparameters were held constant across all ablation conditions to ensure that performance differences reflected feature removal rather than differences in model tuning. The analysis used the same LOGO cross-validation framework and encoding strategies as described previously across nine sST datasets.

#### 1.4. Domain-shift stress testing and cross-platform evaluation

To assess robustness to platform-level differences in score distributions, we perturbed the whole-cell communication score of each held-out dataset within the LOGO-CV framework. We first standardised the whole-cell communication score using the mean ($\mu_x$) and standard deviation ($\sigma_x$) of the held-out dataset:$z = \frac{x-\mu_x}{\sigma_x}$. We applied four perturbations independently:

- **Mean shift:** $z' = z + \Delta\mu$, where $\Delta\mu \in \{-0.5, +0.5\}$.
- **Variance scaling:** $z' = sz$, where $s \in \{0.6, 0.8, 1.25, 1.5\}$.
- **Shape distortion:** $z' = \text{sign}(z)|z|^p$, where $p \in \{0.7, 1.4\}$, preserving rank order.
- **Noise:** $z' = z + \epsilon$, where $\epsilon \sim \mathcal{N}(0, \sigma^2)$ and $\sigma \in \{0.2, 0.5, 1.0\}$. Each noise setting was repeated using five random seeds.

We mapped the perturbed values back to the original scale, $x' = z'\sigma_x + \mu_x$ and passed them to the frozen models. We evaluated predictions against the original cytoplasmic and nuclear scores using $R^2$ and NRMSE. For each dataset, we averaged results across noise repetitions. To assess the generalisability of CCIDeconv across spatial transcriptomics technologies, we evaluated the framework on an external CosMx dataset. We loaded the CosMx data into spatial experiment objects and processed them using the same analytical pipeline as for the 10x Genomics Xenium datasets. We then applied the pretrained CCIDeconv Spatial Procedure (SP) model directly to the processed CosMx data.

##### 1.5. Nuclear compartment characteristics

To investigate factors associated with differences in nuclear and cytoplasmic prediction performance, six dataset-level characteristics were calculated for each of the nine ST datasets:

- Nuclear detected genes: the median number of genes with non-zero expression detected per nucleus/cell.

- Nuclear detection ratio: the ratio of the median number of detected genes in the nuclear compartment to that in the cytoplasmic compartment.

$$\text{Nuclear Detection Ratio} = \frac{\text{Median nuclear detected genes per cell}}{\text{Median cytoplasmic detected genes per cell}}$$

- Nuclear zero-count fraction: the median proportion of genes with zero expression across nuclei/cells, used as a measure of nuclear transcript sparsity.
- Nuclear–cytoplasmic dropout gap: the difference between the nuclear and cytoplasmic zero-count fractions.

$$\text{Dropout Gap} = \text{Median nuclear zero fraction} - \text{Median cytoplasmic zero fraction}$$

- N/C expression ratio: Median gene-wise ratio of total transcript abundance between compartments (for all genes with non-zero cytoplasmic counts):

$$\text{N/C Ratio} = \text{median}\left(\frac{\text{Total nuclear counts per gene}}{\text{Total cytoplasmic counts per gene}}\right)$$

- Total panel gene count: the total number of genes represented in the expression panel for each dataset, providing a measure of available gene coverage.

We extracted the metrics (nuclear and cytoplasmic $R^2$ and NRMSE, as well as the differences between compartments, where $\Delta$ $R^2$ is the cytoplasmic-nuclear $R^2$ and $\Delta$ NRMSE is nuclear-cytoplasmic NRMSE) pertaining to the LOGO CV robustness analysis for SP and ScP procedures of CCIDeconv. We correlated the dataset characteristic with the above mentioned metrics.

##### 1.6. End-to-end evaluation

To assess the effect of classification errors on regression performance, we additionally evaluated CCIDeconv across all events. Events classified as interacting retained their allocated regression scores, whereas events classified as non-interacting were assigned the standardized equivalent of a raw score of zero. We compared regression metrics calculated across all events with those calculated conditionally on events included in regression. Changes in $R^2$ and NRMSE quantified the impact of cascading classification errors. All allocation parameters were estimated using training data only.

#### References

Gabriela Rapozo Guimarães, Giovanna Resk Maklouf, Cristiane Esteves Teixeira, Leandro de Oliveira Santos, Nayara Gusmão Tessarollo, Nayara Evelin de Toledo, Alessandra Freitas Serain, Cristóvão Antunes de Lanna, Marco Antônio Pretti, Jéssica Gonçalves Vieira da Cruz, Marcelo Falchetti, Mylla M Dimas, Igor Salerno Filgueiras, Otavio Cabral-Marques, Rodrigo Nalio Ramos, Fabiane Carvalho de Macedo, Fabiana Resende Rodrigues, Nina Carrossini Bastos, Jesse Lopes da Silva, Edroaldo Lummertz da Rocha, Cláudia Bessa Pereira Chaves, Andreia Cristina de Melo, Pedro M M Moraes-Vieira, Marcelo A Mori, and Mariana Boroni. Single-cell resolution characterization of myeloid-derived cell states with implication in cancer outcome. *Nat. Commun.*, 15(1):5694, July 2024.

Yuhan Hao, Tim Stuart, Madeline H Kowalski, Saket Choudhary, Paul Hoffman, Austin Hartman, Avi Srivastava, Gesmira Molla, Shaista Madad, Carlos Fernandez-Granda, and Rahul Satija. Dictionary learning for integrative, multimodal and scalable single-cell analysis. *Nat. Biotechnol.*, 42(2):293–304, February 2024.

Jie Pan, Jaume Fores-Martos, Claire Delpirou Nouh, Tanner D Jensen, Kristen Vallejo, Romain Cayrol, Saman Ahmadian, Euan A Ashley, Michael D Greicius, and Inma Cobos. Deciphering glial contributions to CSF1R-related disorder via single-nuclear transcriptomic profiling: a case study. *Acta Neuropathol. Commun.*, 12(1):139, August 2024.

Davide Risso and Michael Cole. *scRNAseq: Collection of Public Single-Cell RNA-Seq Datasets*, 2026. URL https://bioconductor.org/packages/scRNAseq. R package version 2.27.0.

Cristian Ruiz-Moreno, Sergio Marco Salas, Erik Samuelsson, Sebastian Brandner, Mariette E G Kranendonk, Mats Nilsson, and Hendrik G Stunnenberg. Harmonized single-cell landscape, intercellular crosstalk and tumor architecture of glioblastoma. *bioRxiv*, August 2022.

Rapolas Zilionis, Camilla Engblom, Christina Pfirschke, Virginia Savova, David Zemmour, Hatice D Saatcioglu, Indira Krishnan, Giorgia Maroni, Claire V Meyerovitz, Clara M Kerwin, Sun Choi, William G Richards, Assunta De Rienzo, Daniel G Tenen, Raphael Bueno, Elena Levantini, Mikael J Pittet, and Allon M Klein. Single-cell transcriptomics of human and mouse lung cancers reveals conserved myeloid populations across individuals and species. *Immunity*, 50(5):1317–1334.e10, May 2019.

#### 2. Supplementary Figures

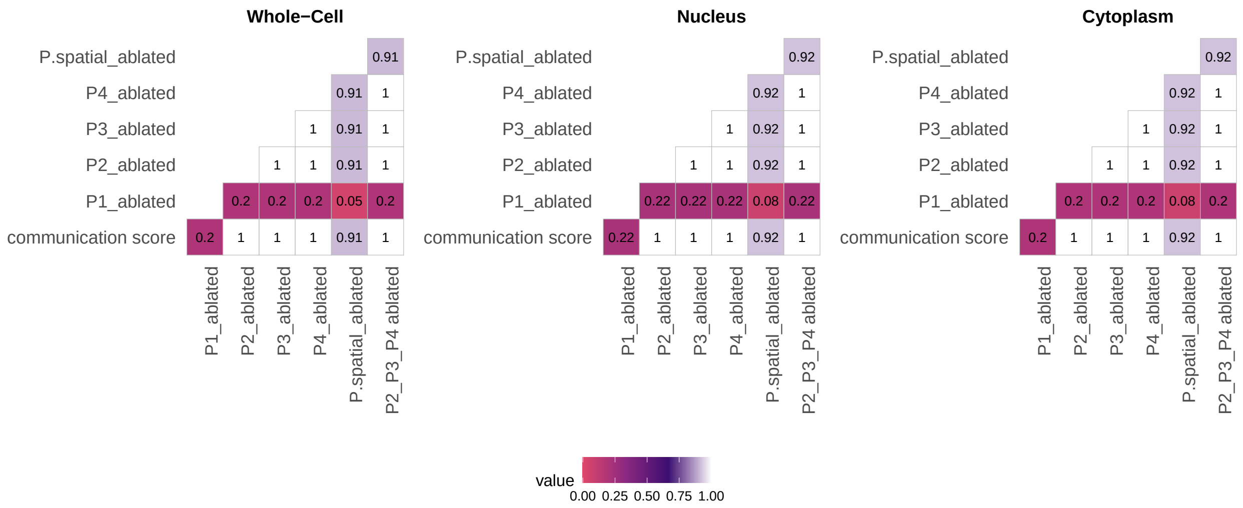


Supplementary Fig. 1: Correlation of the communication score with the various ablated components of the CellChat formula. P1 - Hill function of LR expression, P2 - Antogonist expression, P3 - Agonist expression, P.spatial - spatial distance $(S_i, j)$

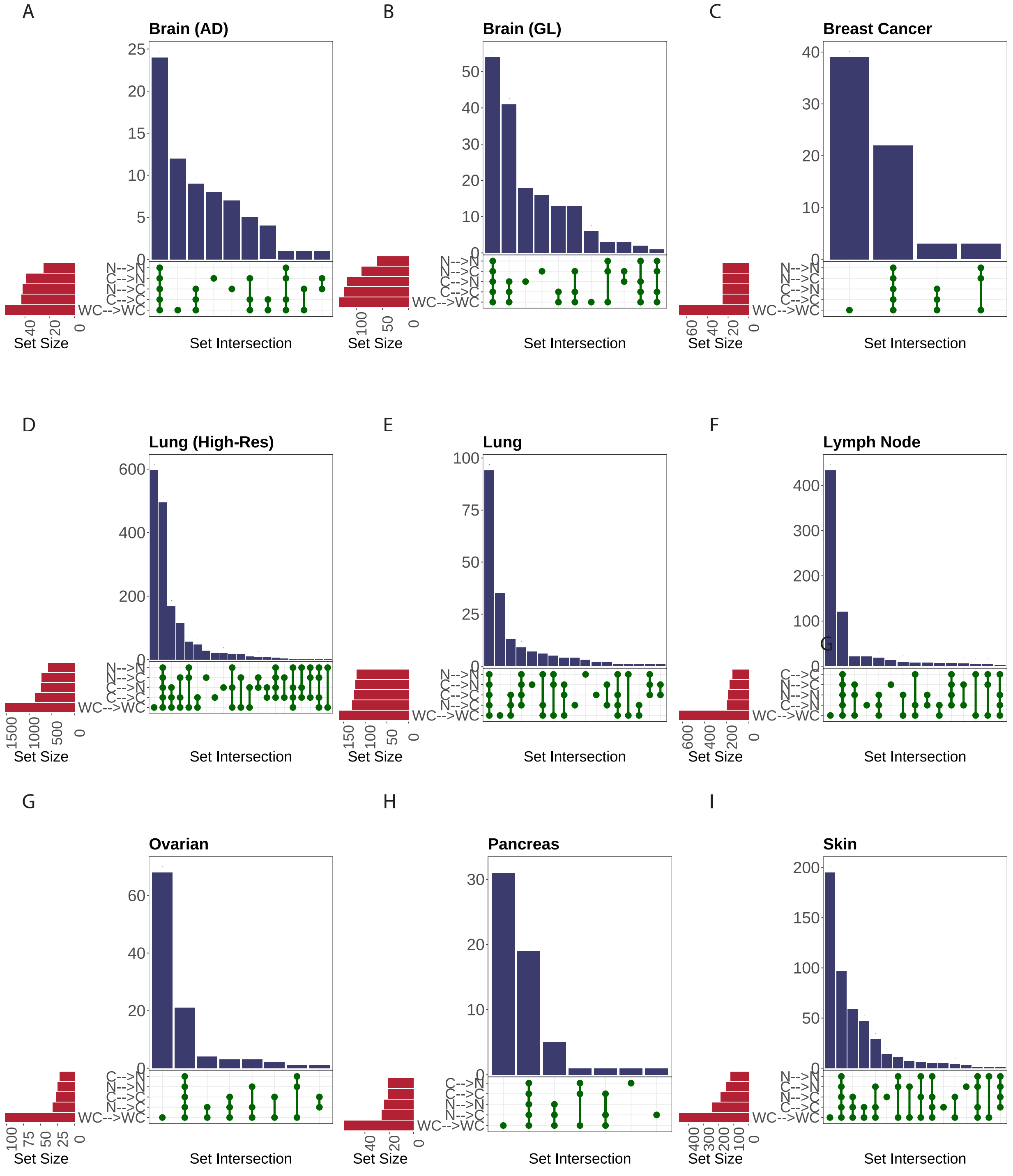


Supplementary Fig. 2: Upset plots of the significant LR communicating between each of the subcellular regions for all the nine subcellularly resolved spatial transcriptomics datasets. A. Brain (AD), B. Brain (GL), C. Breast Cancer, D. Lung (High-Res), E. Lung, F. Lymph Node, G. Pancreas, H. Ovarian, I. Skin. N - Nucleus, C - Cytoplasm, WC - whole cell, AD - Alzheimer's Disease, GL -Glioblastoma.

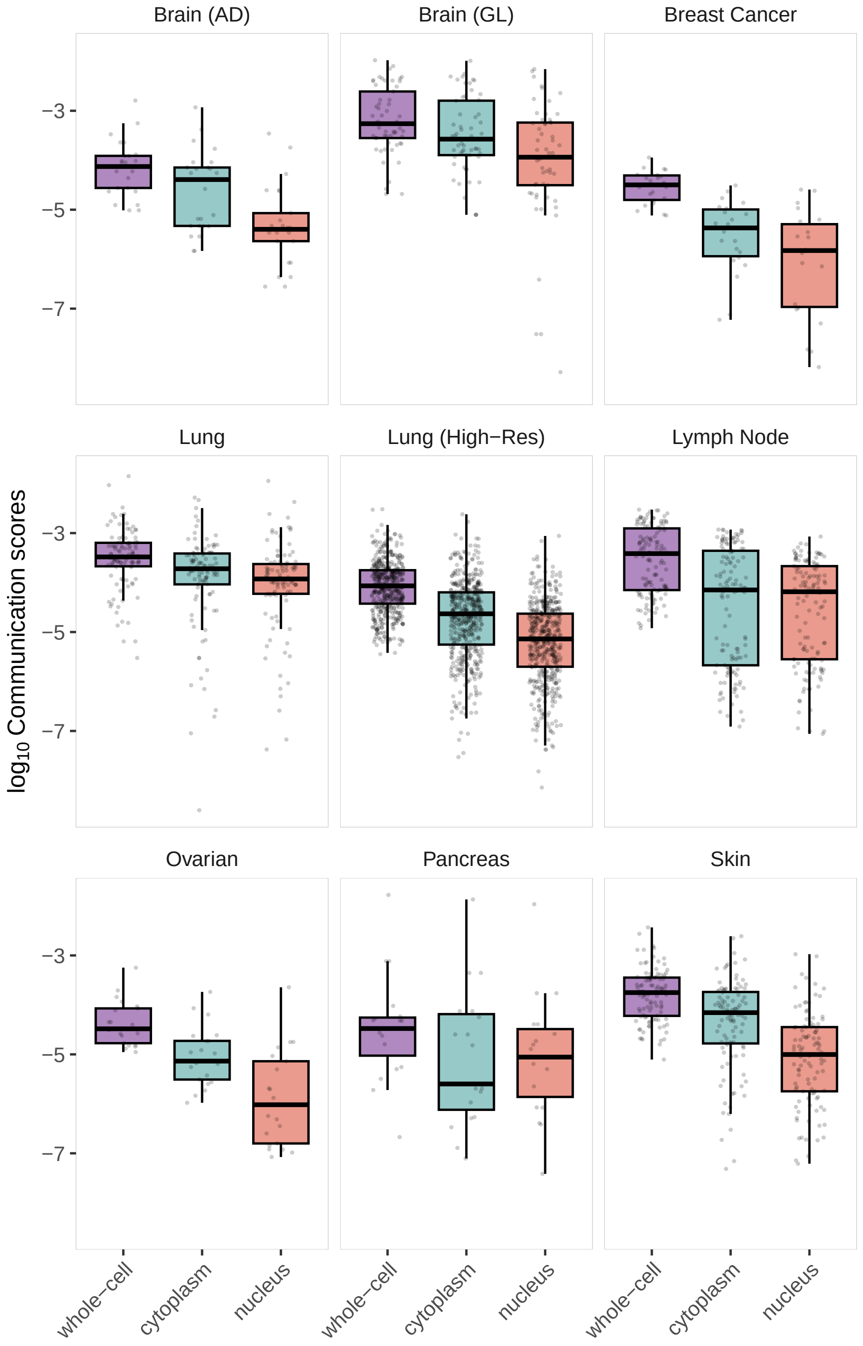


Supplementary Fig. 3: Distribution of communication scores in nine subcellularly resolved spatial transcriptomics datasets across the whole-cell and its subcellular regions. Only the significantly communicating ligand receptor pairs (p-value $< 0.05$) that were identified in all three regions (cell, cytoplasm and nucleus) are shown. AD - Alzheimer's Disease, GL -Glioblastoma.

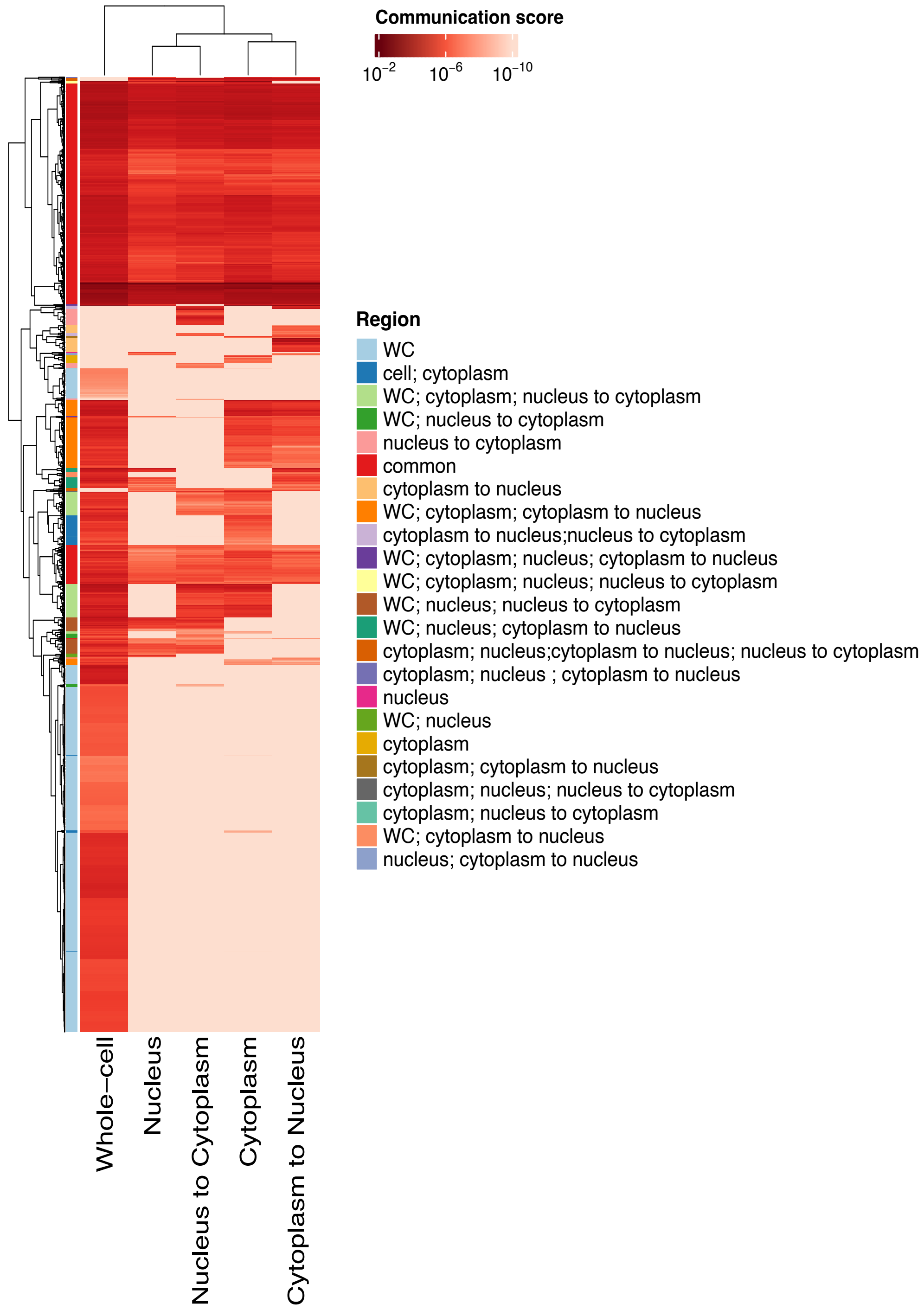


Supplementary Fig. 4: Heatmap of the communication score of significant interactions occurring between the different regions of the cell. WC - Whole-cell

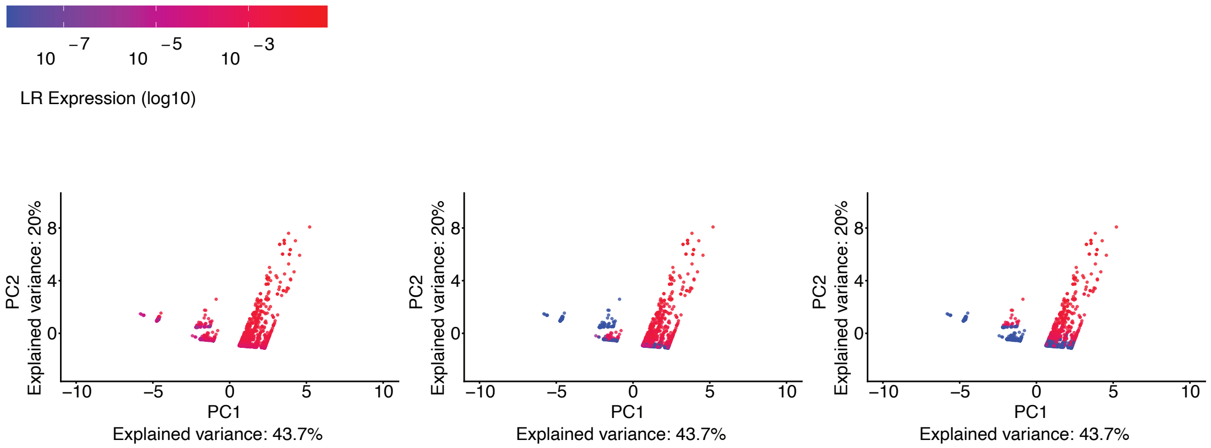


Supplementary Fig. 5: Principal component analysis (PCA) of CellChat-derived communication score components across the whole cell (left), cytoplasm (middle), and nucleus (right) for all the nine subcellularly resolved spatial transcriptomics datasets. Each point represents a ligand–receptor (LR) pair and is colored by the P1 value, defined as the Hill function of LR expression.

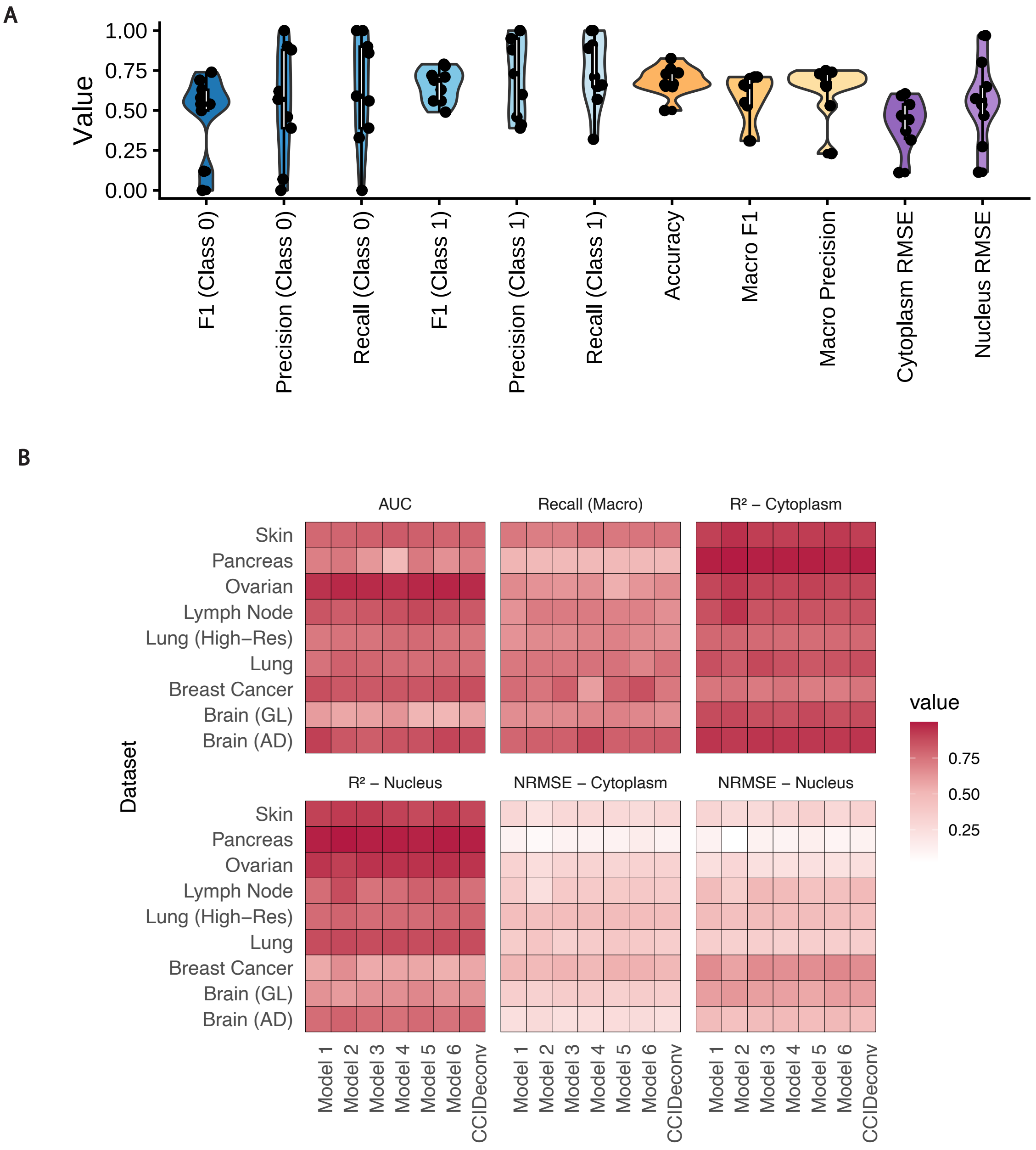


Supplementary Fig. 6: A. Additional performance metrics of CCIDeconv across the nine subcellularly resolved spatial transcriptomics datasets not shown in the main Figure 2. B. Classification and regression metrics of all the models with different base learning configurations and feature encoding strategies of the subcellular location features from CellChatDB and HPA across the nine sST datasets.

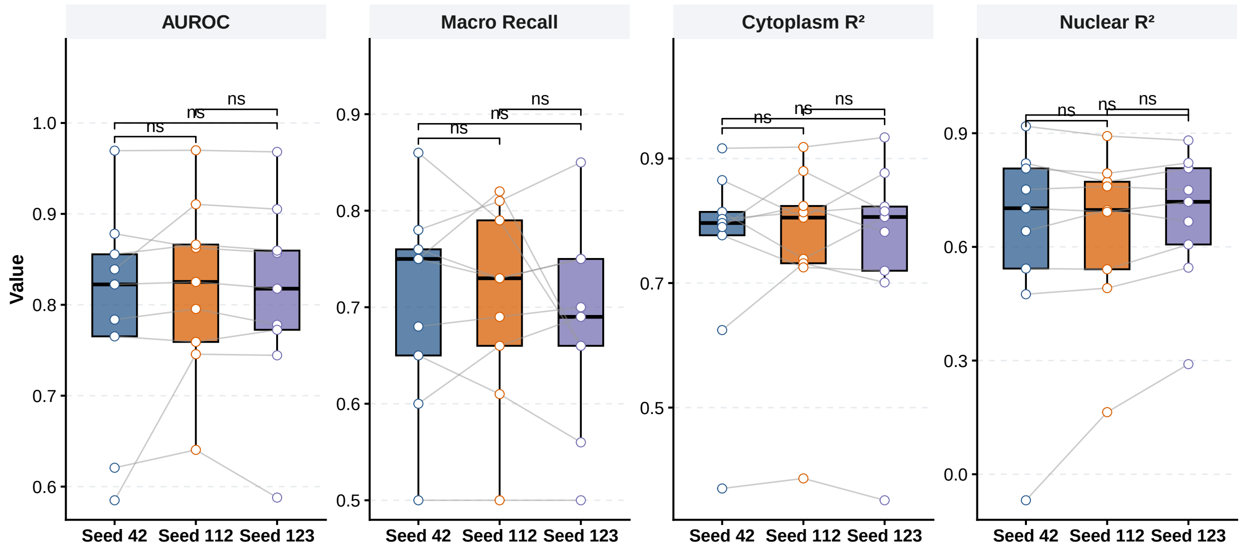


Supplementary Fig. 7: The classification and regression metrics across the 9 datasets evaluated using LOGO -CV on different random seeds. Paired Wilcoxon tests, with P-values adjusted for multiple comparisons using the Benjamini–Hochberg (BH) method was used to compare the performance across the different random seeds.

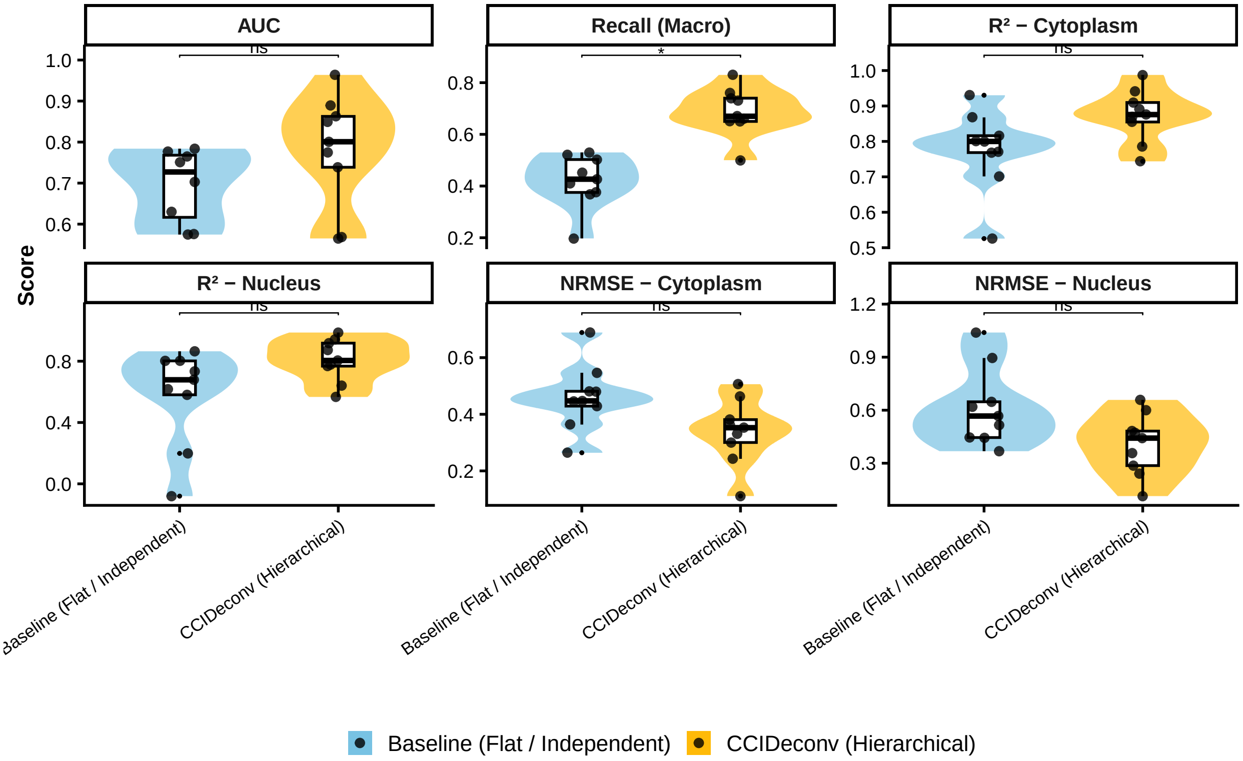


Supplementary Fig. 8: Performance comparison of CCIDeconv against baseline classification and regression models. Violin plots and boxplots show the comparison between the hierarchical CCIDeconv and flat classification / independent regression baseline models across nine datasets. Metrics compared include multiclass OVR AUC, Macro Recall, $R^2$, and NRMSE for both cytoplasm and nucleus compartments. Statistical comparisons between CCIDeconv and each baseline were performed using paired Wilcoxon signed-rank tests, with P-values adjusted for multiple comparisons using the Benjamini–Hochberg (BH) method.

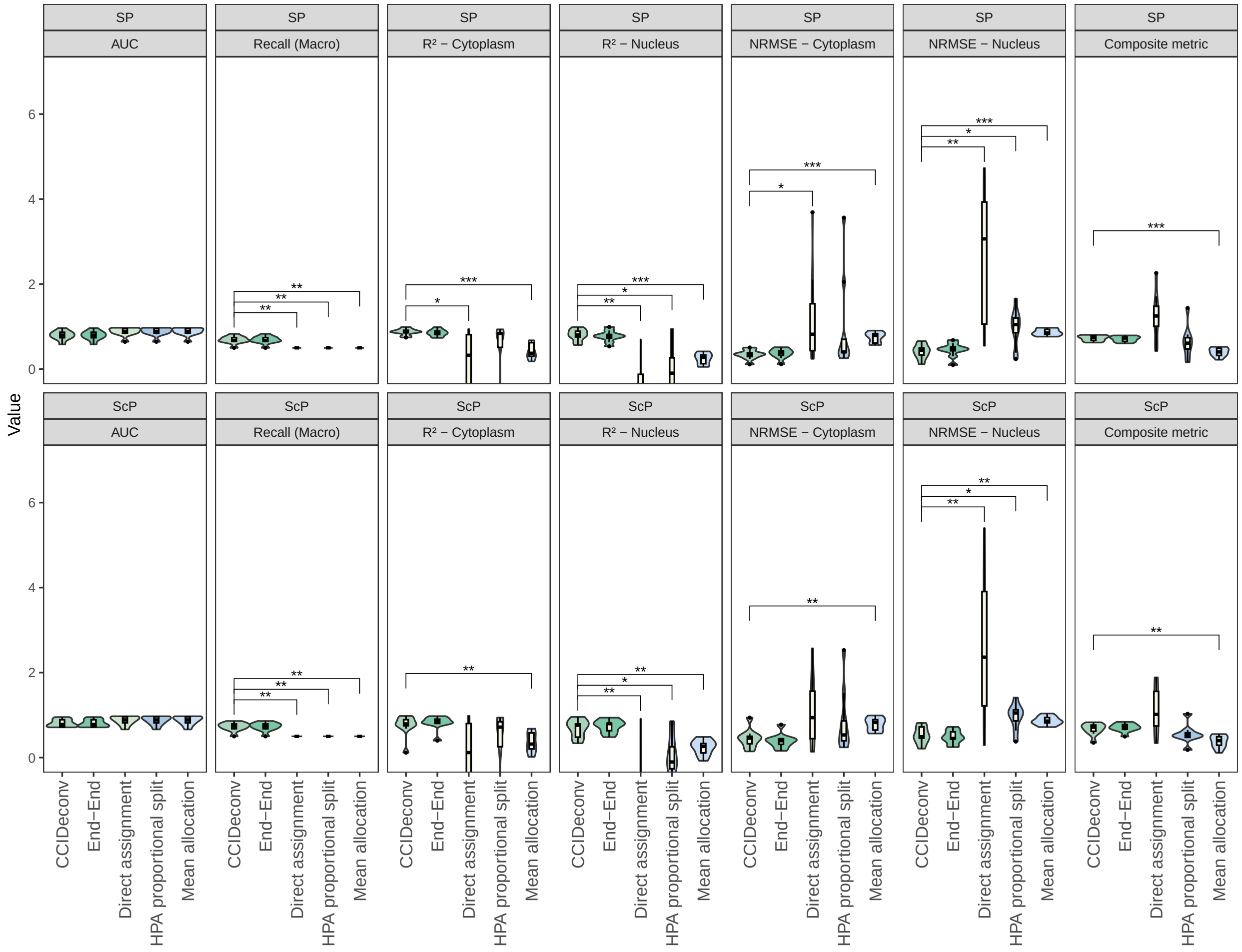


Supplementary Fig. 9: Boxplots show classification and regression performance for direct assignment, HPA proportional split, mean allocation, end-to-end prediction, and CCIDeconv under the SP and ScP procedures. Statistical comparisons between CCIDeconv and each baseline were performed using paired Wilcoxon signed-rank tests, with P-values adjusted for multiple comparisons using the Benjamini–Hochberg (BH) method. Statistically significant differences are indicated in the figure. Classification metrics are identical for end-to-end prediction and CCIDeconv and are therefore shared between these approaches.

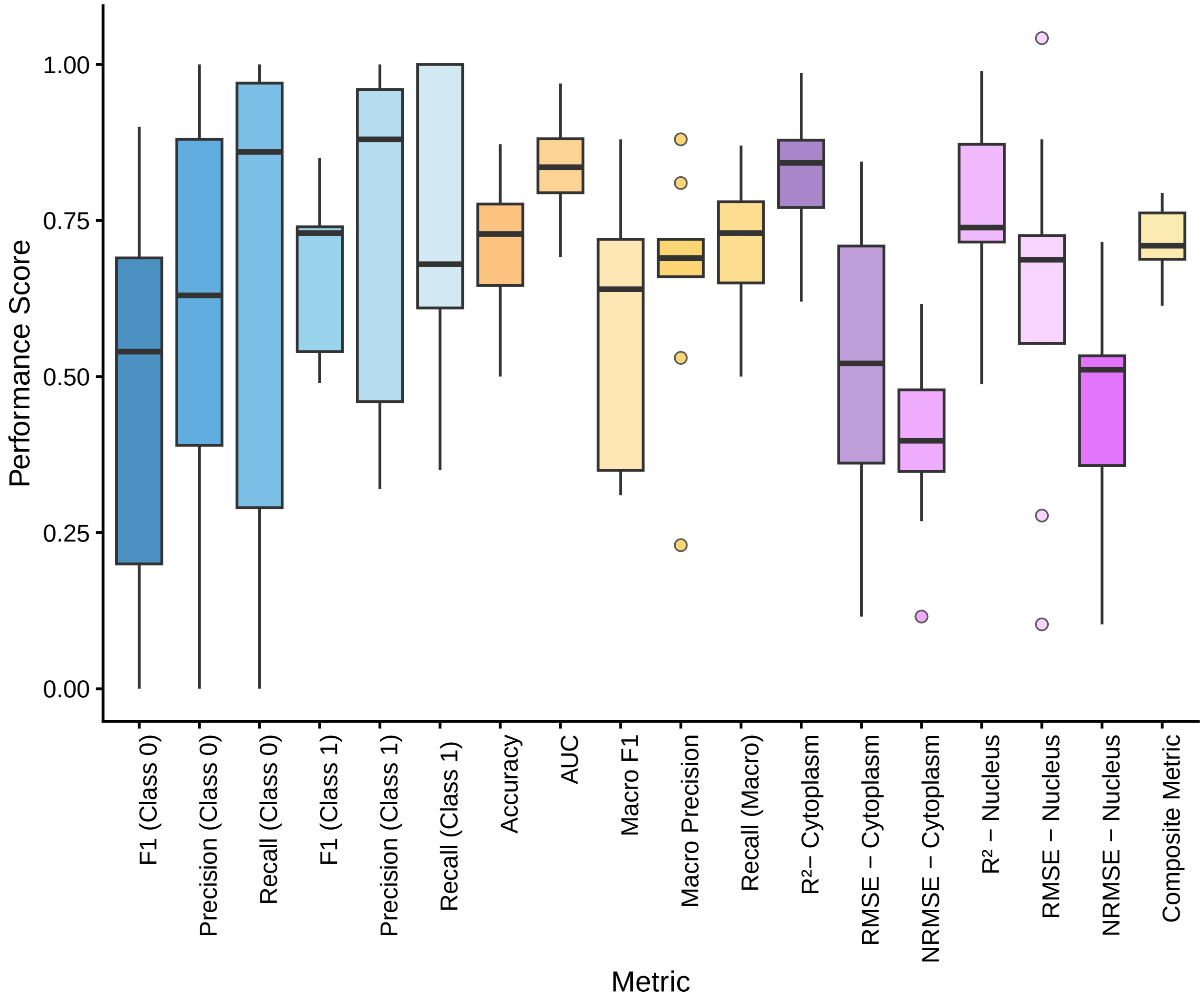


Supplementary Fig. 10: The classification and regression metrics across the 9 datasets evaluated using LOGO -CV on a subset of 3,193 CCI events. These events were significant in the whole-cell only (adjusted P value $\leq$ 0.05)

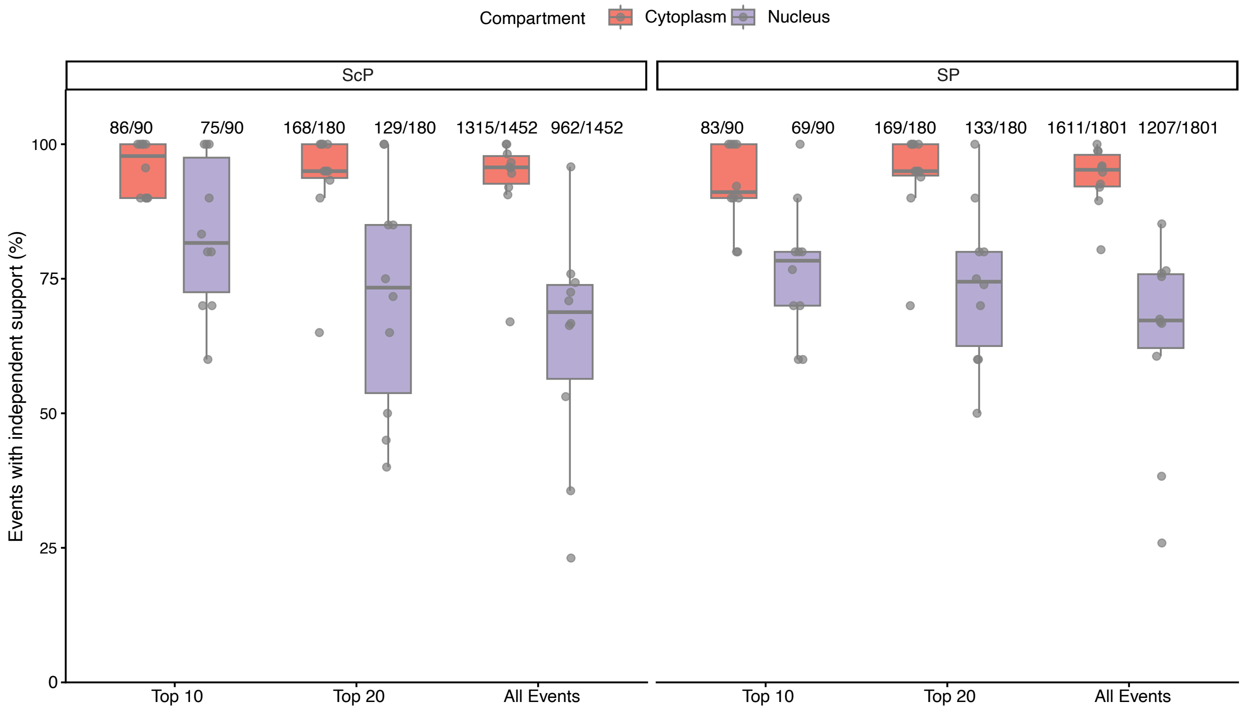


Supplementary Fig. 11: Boxplots of the percentage of LR events with supporting independent evidence for the top 10, 20 and all events, ranked by the predicted cytoplasmic score or nuclear score across the LOGO-CV of the nine datasets for both the procedures of CCIDeconv.

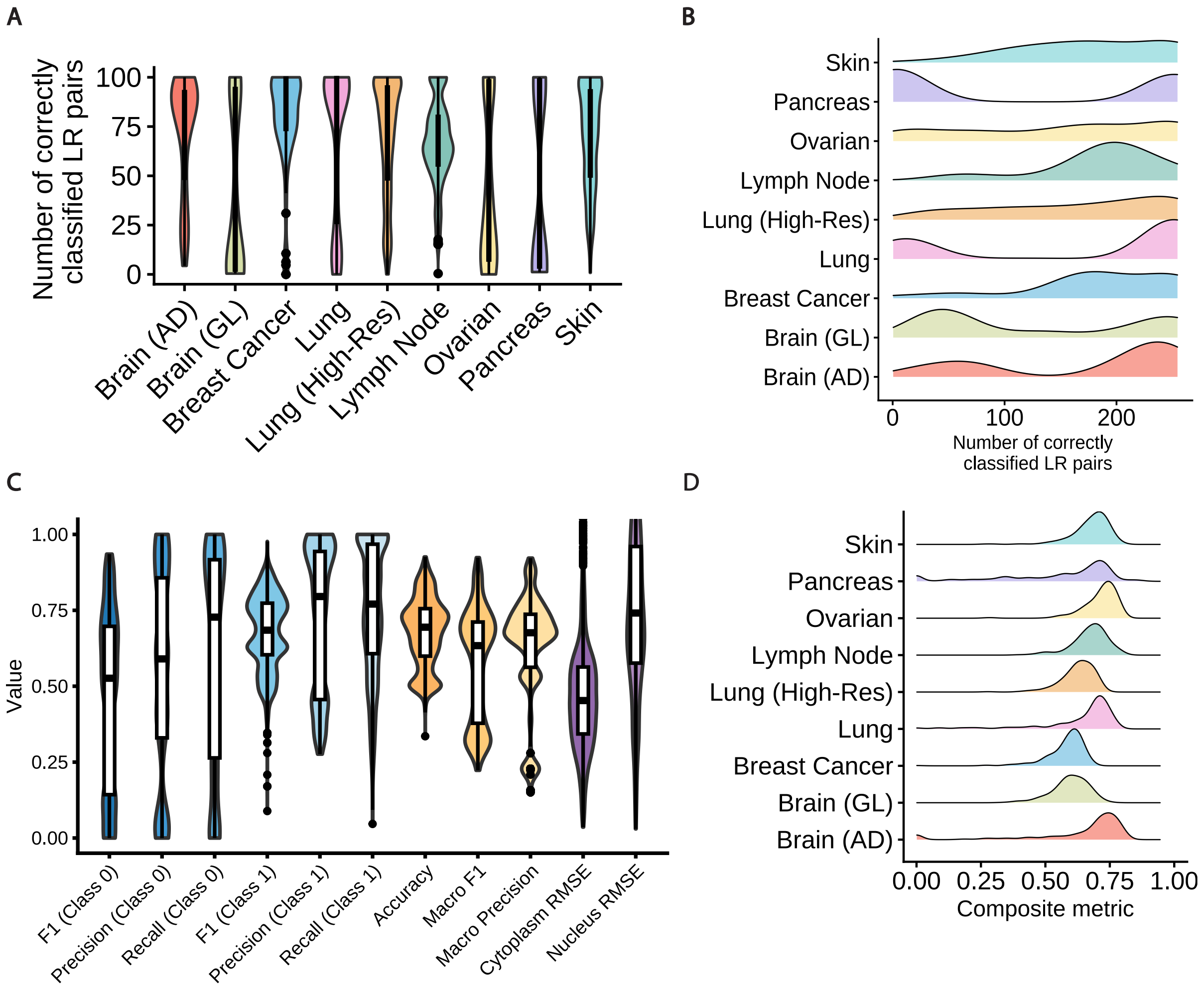


Supplementary Fig. 12: Assessing model stability by training each dataset using varying combinations of the remaining datasets. Each dataset was trained with 255 different combinations of the other eight datasets. A, B. Violin and Ridge plots showing the distribution of the number of correctly classified LR pairs across combinations for each dataset. C. Violin plots of additional performance metrics not reported in the main figure. D. Ridge plots of the composite metric across datasets, illustrating variability when trained on different dataset combinations.

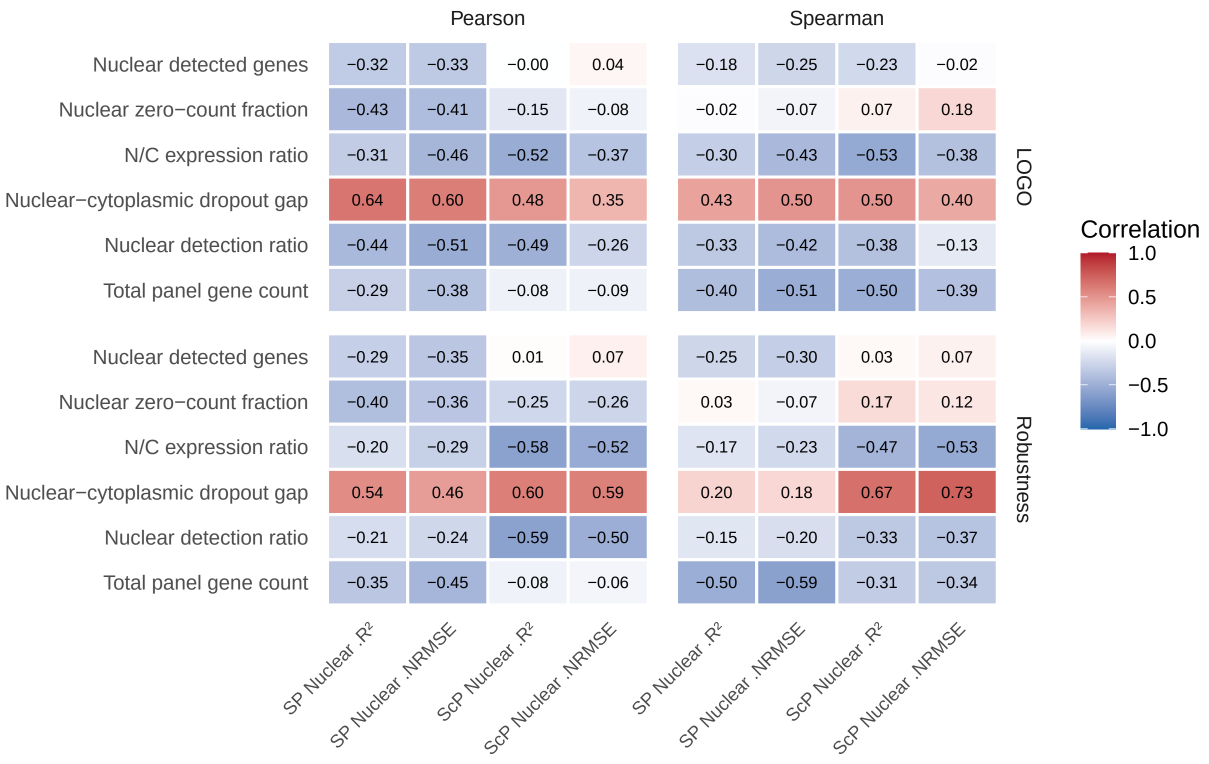


Supplementary Fig. 13: Heatmaps showing Pearson and Spearman correlation coefficients between nuclear dataset characteristics (rows) and compartment-specific prediction performance metrics (columns) under Leave-One-Group-Out (LOGO) and Robustness cross-validation. Comparisons include models trained using SP and ScP. Here, $\Delta R^2$ is defined as cytoplasmic $R^2$ - nuclear $R^2$, and $\Delta$NRMSE as nuclear NRMSE - cytoplasmic NRMSE.

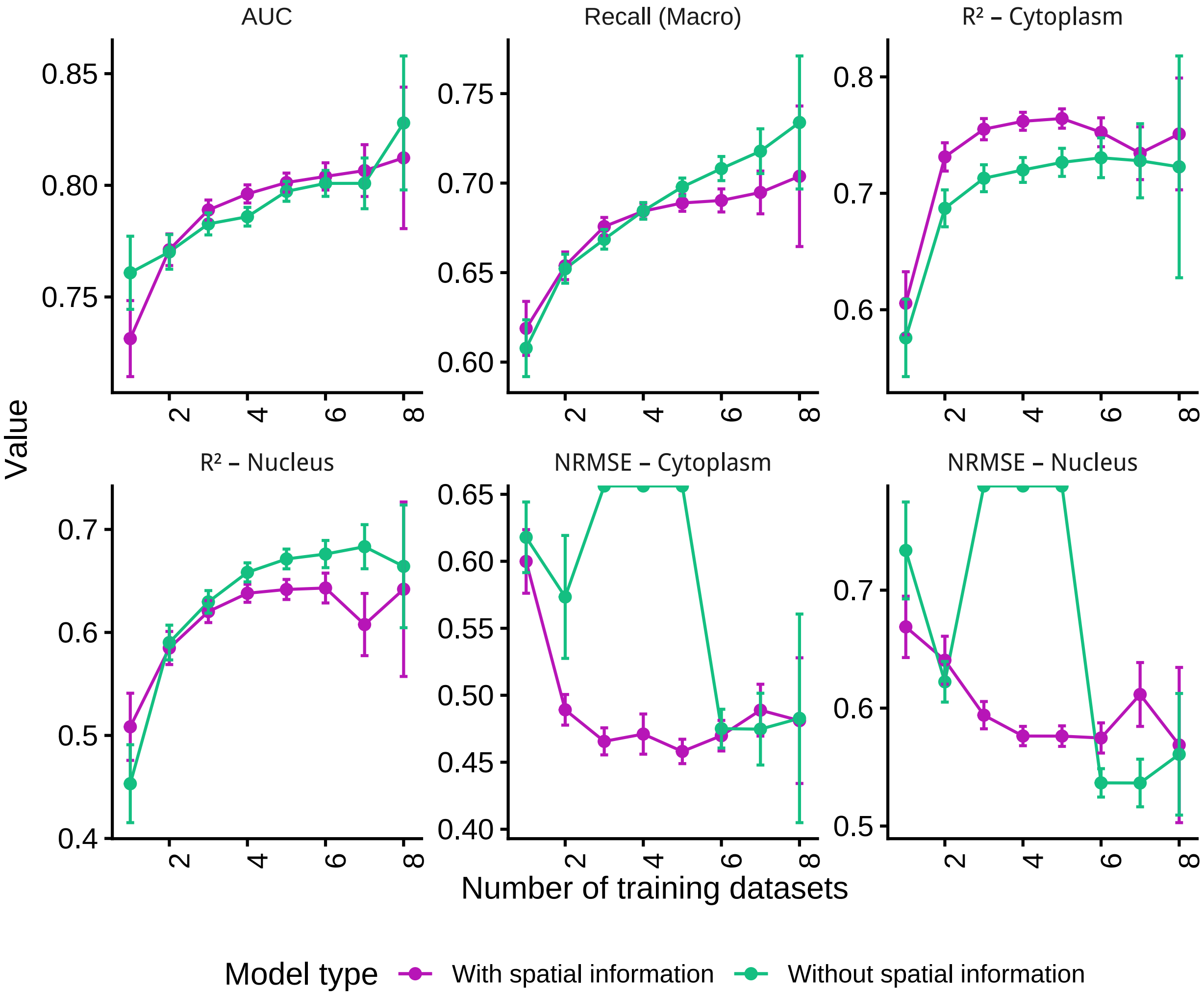


Supplementary Fig. 14: Performance metrics of the models trained with SP and ScP procedures across different combinations of datasets. There were a total of 4590 models trained, with each dataset (n = 9) trained across 255 combinations of other datasets.

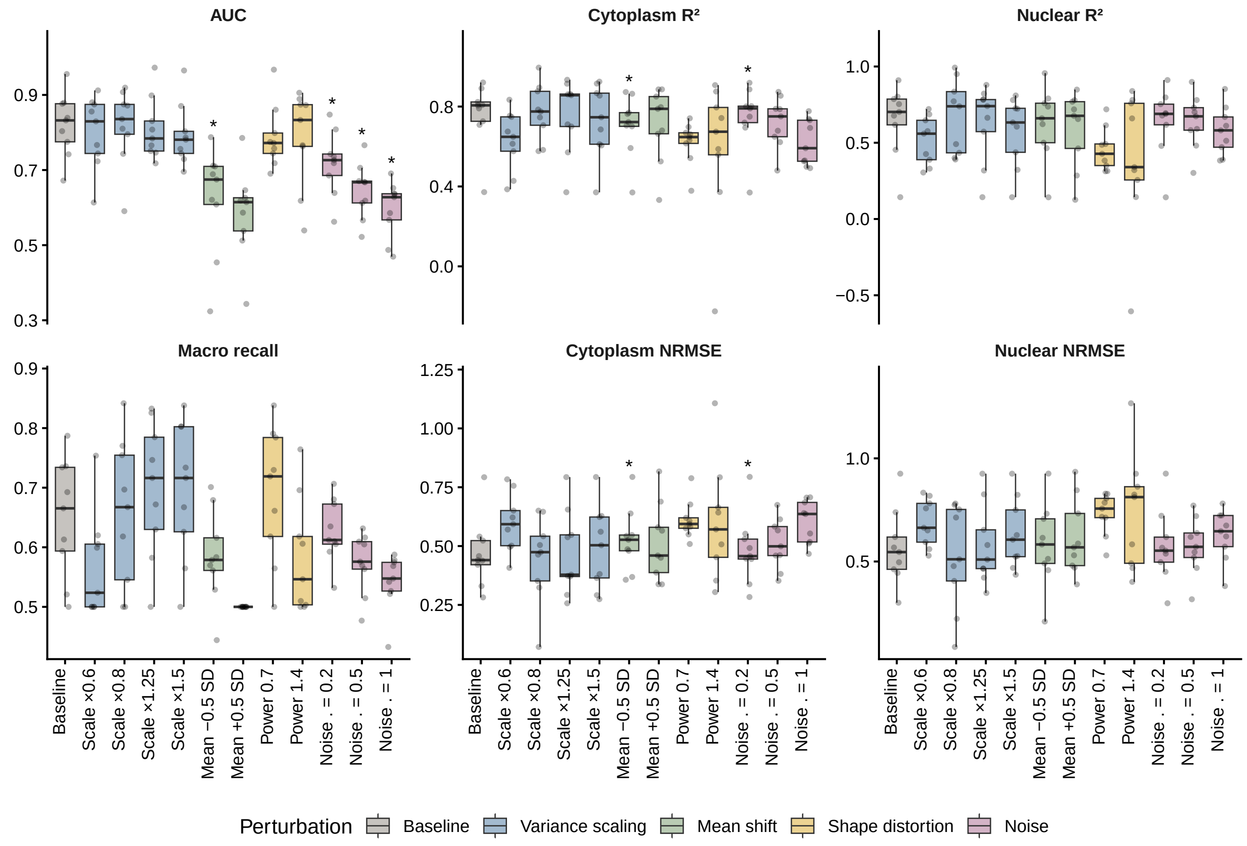


Supplementary Fig. 15: Boxplots of metrics on the SP procedure of CCIDeconv across perturbation types.

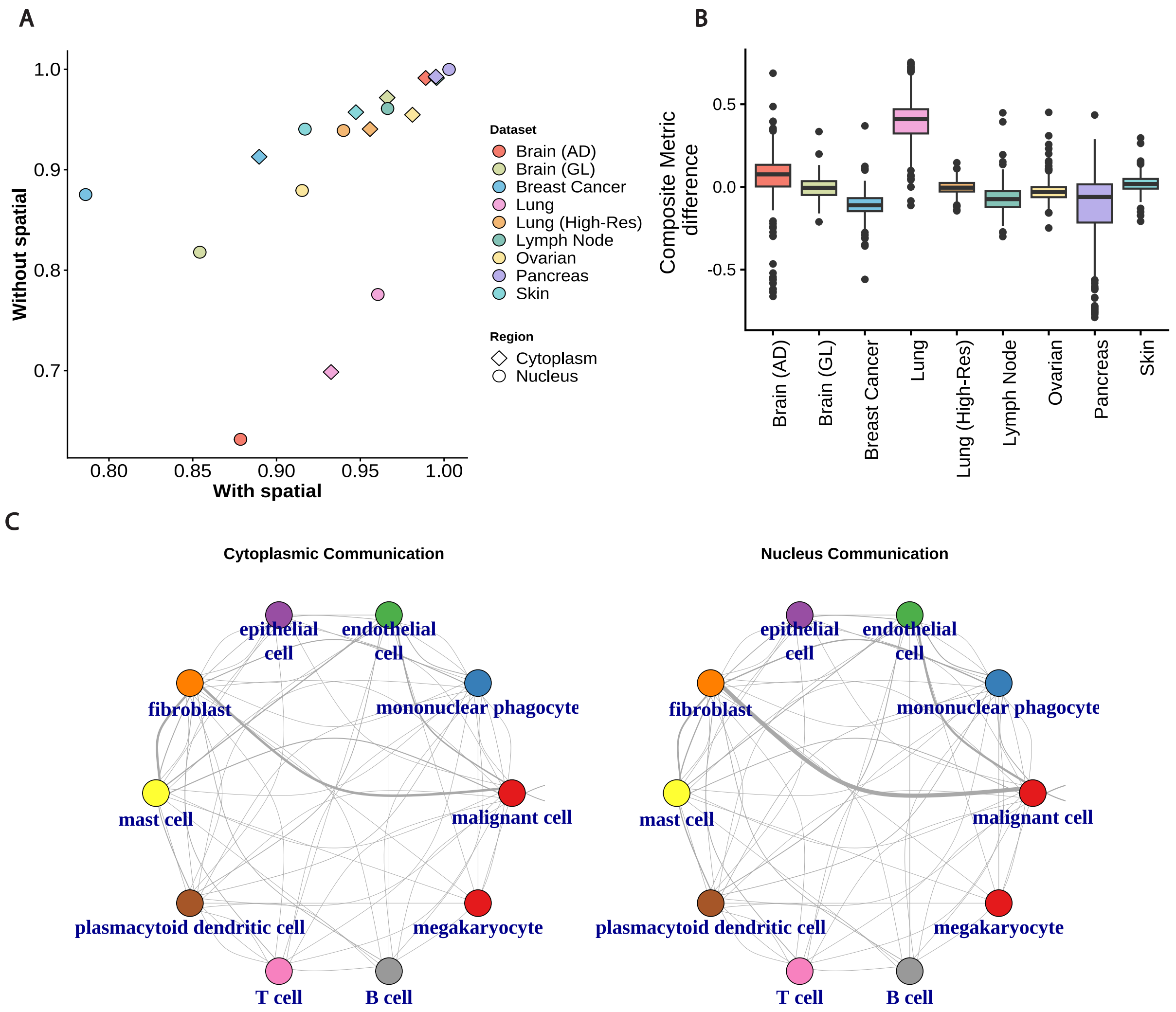


Supplementary Fig. 16: Assessing spatial characteristics of LR communication scores across datasets. A. Correlation of region-specific communication scores with the whole-cell score, comparing models trained with and without spatial features. B. Distribution of pairwise composite metric differences in model predictions across all combinations of training datasets for a given candidate dataset. Brain (AD) and Lung datasets, which exhibit high spatial correlation, are included. C. Network plot showing the total ligand–receptor (LR) scores in the predicted cytoplasm and nucleus when the model is applied to single-cell data.
